# Gadd45 regulates fate decisions of myeloid-type blood cells in *Drosophila*

**DOI:** 10.64898/2025.12.16.694615

**Authors:** Priyasi Jaiswal, Bama Charan Mondal

## Abstract

Growth arrest and DNA damage-inducible 45 (Gadd45) is a conserved, stress-responsive regulator linking cellular stress and cytokine signaling to cell-cycle control, DNA repair, apoptosis, and differentiation. Altered Gadd45 expression has been associated with mammalian solid tumors and hematopoietic malignancies. Gadd45 also supports cytokine-induced myeloid differentiation, though the underlying mechanisms remain unclear. Using the *Drosophila* hematopoietic system and in vivo genetic and cell-biological approaches, we investigated Gadd45’s role in myeloid-type blood cell development. We show that Gadd45 selectively regulates crystal cells by reducing their differentiation and impairing maintenance, while leaving plasmatocyte differentiation unchanged. However, Gadd45 overexpression promotes lamellocyte differentiation, a response typically observed during larval immune challenges and injury. Mechanistically, Gadd45 overexpression induces lamellocyte differentiation through Toll, JNK, and JAK/STAT signaling. Furthermore, Gadd45 overexpression does not induce cell death but alters cell-cycle status in the lymph gland. Our study provides new insight into Gadd45’s specific role in crystal cell development and lamellocyte-associated differentiation in *Drosophila*.

**Significance Statement:**

- Loss of Gadd45 causes a reduction in crystal cell differentiation and impairs their maintenance, while plasmatocyte differentiation remains unchanged.
- Gadd45 overexpression causes myeloid-type progenitor cells to differentiate into lamellocytes, as well as plasmatocytes to transdifferentiate into lamellocytes. Gadd45 triggers differentiation by activating a combination of Toll, JNK, and JAK/STAT signaling pathways.
- Overexpressing Gadd45 does not induce apoptosis but alters the cell cycle behavior of hematopoietic progenitor cells.

## Introduction

The evolutionarily conserved *Gadd45* (Growth Arrest and DNA Damage-inducible 45) gene family encodes an 18 kDa acidic protein that was identified as a gene whose mRNA is rapidly induced by various DNA damage-inducing agents (Fornace et al., 1992; Papathanasiou et al., 1991). The DNA damage–induced transcription of the *Gadd45* gene is mediated by both p53-dependent and -independent mechanisms (Takekawa & Saito, 1998). The mammalian genome encodes three highly homologous Gadd45 proteins: Gadd45α, Gadd45β, and Gadd45γ (Yang et al., 2013). Gadd45 proteins interact with many other cellular proteins that influence the cell cycle and responses to various cellular stresses, including the JNK-MAP kinase pathway (Hoffman & Liebermann, 2009).

Gadd45 expression is altered in solid tumors and hematopoietic malignancies (Patel et al., 2022). *Gadd45* is also induced by various cytokines during myeloid differentiation, although the molecular mechanism remains unclear. Gadd45 deficiency may alter the behavior of hematopoietic stem and progenitor cells during physiological stress, highlighting the need for further research to understand the long-term effects of *Gadd45* misexpression on myelopoiesis and hematopoietic function. Earlier studies have also highlighted Gadd45’s potential role in innate immune function, an area that is still largely unexplored. Understanding the diverse roles of Gadd45 can be helpful in developing treatments for leukemia and other blood disorders (Gupta et al., 2006; Patel et al., 2022).

Genetic manipulation of Gadd45 in several model systems has revealed its unique and shared roles in development and homeostasis. *Drosophila melanogaster* has a single Gadd45 gene, CG11086 (FlyBase ID: FBgn0033153), which shares 31% identity with human Gadd45β. This makes it a genetically tractable model to study Gadd45-mediated pathways compared to mammals with three homologs (Patel et al., 2022). *Drosophila* Gadd45 is conditionally essential: It is dispensable for baseline viability or development but is required for tissue homeostasis under challenge. It is also upregulated during immune responses and injury but not in response to other stress stimuli, such as genotoxic stress (Peretz et al., 2007; Weavers et al., 2019). Gadd45 expression levels and duration also influence phenotypes: transient overexpression does not cause cell death, whereas prolonged overexpression causes significant apoptosis, affects wing development, impairs imaginal disc regeneration, and disrupts polarity via MAPK-JNK signaling (Camilleri-Robles et al., 2019; Peretz et al., 2007). Tissue-specific effects include lifespan extension and enhanced DNA repair in neuroblasts, likely by reducing neurodegeneration (Plyusnina et al., 2011; Bgatova et al., 2015), whereas overexpression in the female germline affects dorso-ventral polarity, causing eggshell defects and disrupting anterior-posterior localization (Peretz et al., 2007).

*Drosophila* is an excellent model organism with versatile genetic tools, offering insights into its myeloid-type hematopoietic system. The regulatory factors of *Drosophila* hematopoiesis are conserved across vertebrates (Banerjee et al., 2019; Gold & Bruckner, 2014; Sinenko, 2024; Yoon et al., 2023). We used robust *Drosophila* genetics and its larval hematopoietic organ, the lymph gland (**Figure 1A**), to investigate the role of Gadd45 during myeloid-type blood cell development. *Drosophila* has three types of terminally differentiated myeloid-type blood cells, or hemocytes: plasmatocytes, which are akin to mammalian macrophages and make up ∼95% of the total blood cells; they secrete antimicrobial peptides and eliminate debris through phagocytosis (Brooks et al., 2023; Maurya et al., 2024; Siekhaus & Stanley-Ahmed, 2024). Crystal cells play a crucial role in melanization, wound healing, blood clotting, and oxygen transport (Bidla et al., 2007; Shin et al., 2024). Signaling pathway defects that maintain progenitors or immune challenges can drive differentiation into larger, specialized cells known as lamellocytes (Banerjee et al., 2019; Kharrat et al., 2025). These cells primarily encapsulate larger invaders, such as parasitic wasp eggs, that exceed plasmatocytes’ engulfment capacity. Since lamellocytes are typically absent under normal, uninduced conditions, their presence in naive animals signifies a disruption in the mechanisms that maintain hemocyte progenitor homeostasis during larval development (Banerjee et al., 2019; Kharrat et al., 2025). The *Drosophila* larval lymph gland contains a supportive niche, stem-like progenitor cells, intermediate progenitors, and mature blood cells (**Figure 1A**), similar to vertebrate hematopoietic organs. Studies from various laboratories, including ours, examined the signaling mechanisms that regulate the maintenance and differentiation of lymph gland blood progenitors (Banerjee et al., 2019; Jung et al., 2005; Kharrat et al., 2022; Lan et al., 2020; Mondal et al., 2014; Shim et al., 2013). The progenitors of third-instar larval stage cells in the lymph glands exhibit a prolonged G2 phase of the cell cycle (Cho et al., 2024; Goins et al., 2024; Kapoor et al., 2022; Sharma et al., 2019). Single-cell transcriptomic data from the lymph gland indicated variation in Gadd45 expression in different cell types (Cho et al., 2020; Girard et al., 2021). Given Gadd45’s established role in cellular stress responses, we hypothesized that Gadd45-dependent signaling in *Drosophila* hemocytes directly shapes hemocyte fate in a context-dependent manner by promoting or restraining measurable changes in lineage specification, hemocyte maintenance, and progenitor-to-differentiated cell transitions.

**Figure 1:**
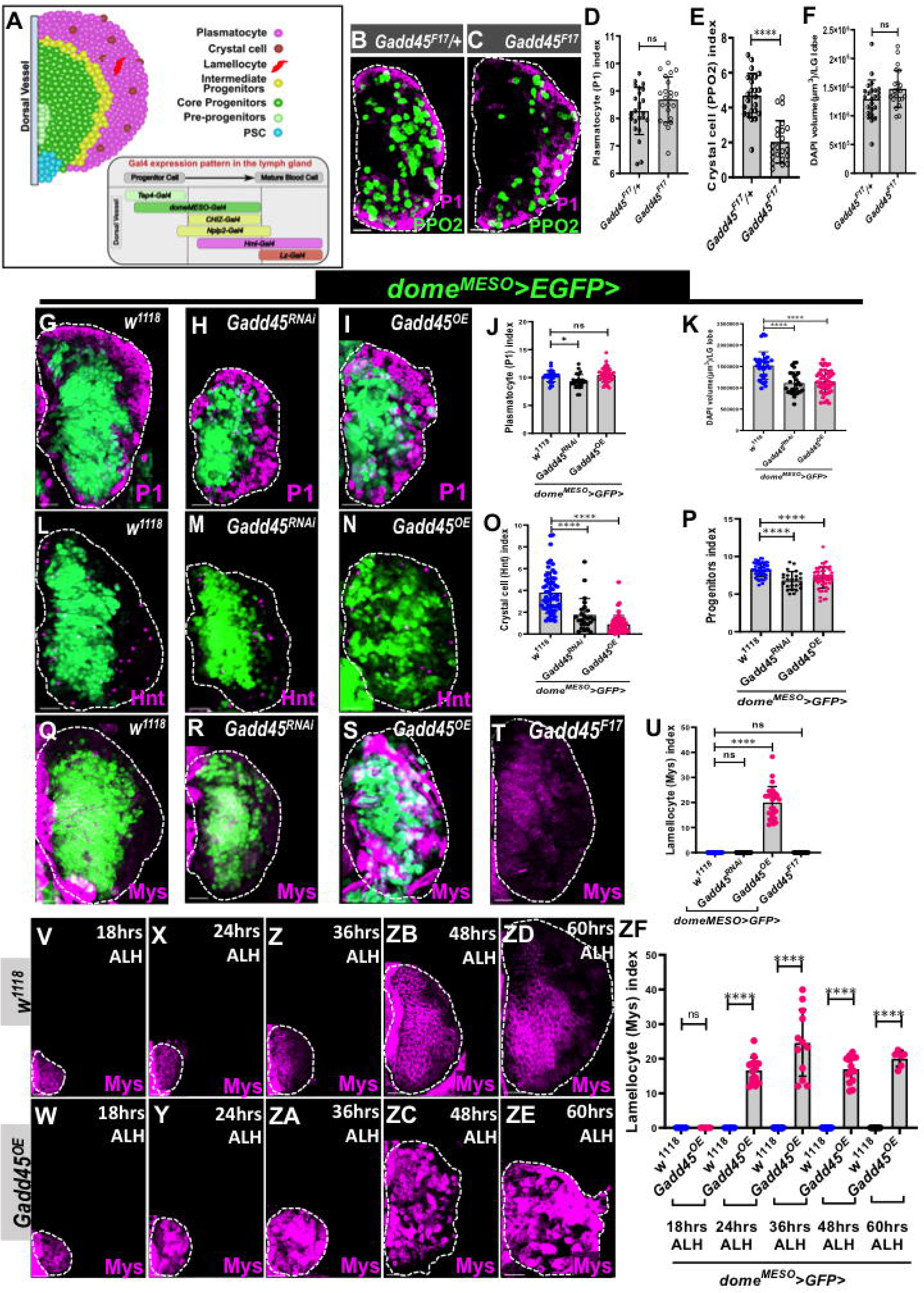
Gadd45 dysregulation in lymph gland progenitors causes lamellocyte differentiation. **(A)** Schematic depicting the distribution of various cell types in the third-instar primary lymph gland lobe and Gal4 drivers marking different stages of blood cell maturation in the lymph gland. **(B-C)** Heterozygous *Gadd45^F17^/+* (B, n=24) and homozygous *Gadd45^F17^* (C, n=22) mutants have similar plasmatocyte (magenta) index whereas, mature crystal cells marked by PPO2 (green) decreases in homozygous *Gadd45^F17^* mutant. **(D)** Quantification of plasmatocyte index in (B-C). **(E)** Quantification of crystal cell index in (B-C). **(F)** Quantification of lymph gland size using DAPI volume in (B-C). **(G-I)** Knockdown of Gadd45 by expressing *UAS-Gadd45^RNAi^*(H, n=27) and continuous expression of Gadd45 by *UAS-Gadd45^OE^* (I, n=46) in lymph gland progenitors (green) using *dome^MESO^-Gal4, UAS-2xEGFP* driver, does not change the number of mature plasmatocytes marked by P1 (magenta) in comparison to control lymph glands (*dome^MESO^-Gal4, UAS-2xEGFP/+*) (G, n=31). **(J)** Quantification of plasmatocyte index in (G-I). **(K)** Quantification of lymph gland size using DAPI volume in (G-I). **(L-N)** Using *dome^MESO^-Gal4, UAS-2xEGFP* driver (green) expression of *UAS-Gadd45^RNAi^* (M, n=24) and *UAS-Gadd4^OE^* (N, n=53) drastically decreases crystal cells, marked by Hnt (magenta), compared to control *w^1118^* (L, n=58) lymph glands. **(O)** Quantification of crystal cell index in (L-N). **(P)** Quantification of progenitor index in (L-N). **(Q-T)** Knockdown of Gadd45 in lymph gland progenitors (green) using the *dome^MESO^-Gal4, UAS-2xEGFP* driver by expressing *Gadd45^RNAi^* (R, n=40) and *Gadd45^F17^* (T, n=18) mutants does not induce lamellocyte differentiation in the lymph gland. In contrast, expressing *Gadd4^OE^* (S, n=28) in progenitors significantly increases lamellocytes marked by Mys (magenta) compared with control *w^1118^* (Q, n=30) lymph glands. **(U)** Quantification of lamellocytes index in (Q-T). **(V-ZE)** Expression of *Gadd45* in lymph gland progenitors using the *dome^MESO^-Gal4* driver does not alter Mys expression at 18 h ALH (W, n = 4), comparable to control lymph glands (*dome^MESO^-Gal4/+*) (V, n = 4). At 24 h ALH, *Gadd45* overexpression initiates lamellocyte differentiation (Y, n = 14), which progressively increases at 36 h ALH (ZA, n = 12) and results in extensive lamellocyte production by 48 h ALH (ZC, n = 12). In contrast, control lymph glands at 24 h (X, n = 12), 36 h (Z, n = 10), and 48 h ALH (ZB, n = 10) retain Mys expression primarily in progenitor cells. By 60 h ALH, control lymph glands continue to exhibit progenitor-level Mys expression (ZD, n=6), whereas *Gadd45*-overexpressing lymph glands show strong Mys upregulation, consistent with extensive lamellocyte differentiation and disintegration of the cortical zone (ZE, n=6). **(ZF)** Quantification of lamellocytes index in (V-ZE). All confocal images represent maximum-intensity projections of the middle third optical sections of the wandering third instar lymph gland lobe from at least three independent biological experiments. A lymph gland lobe is outlined by white dashed lines. Scale bar represents 25 μm. Error bars in graphs: mean ± S.D. ns.: not significant (p>0.01); ****p<0.00001. n represents lymph gland lobe numbers.

Here, we investigated the functions of Gadd45 during *Drosophila* hematopoiesis. We show that the absence of Gadd45 reduces the number of crystal cells, although plasmatocyte differentiation remains largely unaffected. Conversely, Gadd45 overexpression prompts intermediate progenitors to differentiate into lamellocytes, and plasmatocytes to transdifferentiate into lamellocytes—a response typically observed during the larval stage only in reaction to immune challenges and injury. Interestingly, our results further show that Gadd45 overexpression triggers a differentiation program by activating the combined Toll, JNK, and JAK/STAT signaling pathways, as reported during the injury-induced inflammatory response (Evans et al., 2022).

## Results

### Gadd45 dysregulation in lymph gland progenitors caused lamellocyte differentiation

We first examined third-instar larval lymph glands in a Gadd45 null mutant, *Gadd45^F17^* (Nelson et al., 2016). The third-instar larval lymph gland size, differentiation of plasmatocytes marked by P1 (also known as NimC1) (Kurucz et al., 2007), and lamellocytes marked by Myospheroid (Mys, a βPS integrin)(Evans et al., 2022; Madhwal et al., 2020) in *Gadd45* null mutants were similar to those in the controls (**Figures 1B-D, F, Q, T, U**). However, in mutant lymph glands, the number of crystal cells decreased significantly, as indicated by prophenoloxidase 2 (PPO2) immunostaining (Shin et al., 2024) (**Figures 1B-C, E**). We also confirmed the genomic deletion of the coding portion of the Gadd45 gene in the *Gadd45^F17^*(BL81039) mutant by PCR on genomic DNA (**Figure S1A**), as reported in the original article (Nelson et al., 2016).

To dissect the spatial requirement for Gadd45 within the lymph gland, we used multiple *Gal4* drivers with distinct expression domains in the larval lymph gland (**Figure 1A**)(Kharrat et al., 2022). *dome^MESO^-Gal4* is expressed in most medullary zone progenitors, including both core and distal populations (Girard et al., 2021; Spratford et al., 2021), whereas *Tep4-Gal4* is restricted to core progenitors that are quiescent and less responsive to immune challenge (Girard et al., 2021). *CHIZ-Gal4* marks a transitional intermediate progenitor population primed for differentiation (Spratford et al., 2021), whereas *Nplp2-Gal4* is expressed in the intermediate progenitor zone (Cho et al., 2020). *Hml*^Δ^*-Gal4* expression begins in intermediate progenitors, with higher expression in differentiated hemocytes (Girard et al., 2021; Spratford et al., 2021), and *Lz-Gal4* is expressed only in crystal cells (Koranteng et al., 2020; Mukherjee et al., 2011).

We then depleted Gadd45 using a pre-validated RNA interference (RNAi) transgenic fly line (Burbridge et al., 2021; Weavers et al., 2019; Xu et al., 2022). Quantitative RT-PCR analysis of the third-instar lymph gland also revealed a significant reduction in Gadd45 transcript levels (**Figure S1B**). Gadd45 depletion with the *dome^MESO^-Gal4* driver (*dome^MESO^*>*GFP*>*Gadd45^RNAi^*) did not cause a highly significant change in lymph glands’ differentiation pattern of plasmatocytes (**Figures 1G-H, J**) and lamellocytes (**Figures 1Q-R, U**) compared to the controls (*dome^MESO^>GFP/+*). However, downregulating Gadd45 resulted in smaller lymph glands with a decrease in progenitor population (**Figures 1K, P**), and the number of crystal cells marked by the transcription factor Hindsight (Hnt, also known as pebbled) (Rai et al., 2026) decreased significantly compared to the control (**Figures 1L-M, O**).

Next, we used pre-validated *UAS-Gadd45* lines to overexpress Gadd45 in the progenitor population and determine whether this caused any phenotype, as reported earlier in several tissues (Camilleri-Robles et al., 2019; Peretz et al., 2007; Weavers et al., 2019). Remarkably, *dome^MESO^-Gal4*-mediated Gadd45 overexpression caused a significant increase in the number of lamellocytes (**Figures 1Q, S, and U**), along with premature dispersal of the lymph gland primary lobes leading to a decrease in lymph gland size, progenitor index (**Figures 1K, P**), and crystal cell (Hnt) index (**Figures 1L, N, and O**); however, there was no significant change in plasmatocytes (**Figures 1G, I, and J**).

The progenitor driver *dome^MESO^-Gal4* is expressed in the early first-instar larval lymph gland (**Figure S1C**), and Gadd45 overexpression disrupts lymph gland architecture during the wandering third-instar stage and promotes lamellocyte differentiation (**Figure 1S**). To study this developmental process and the temporal expression when lamellocytes begin to differentiate upon ectopic Gadd45 expression, we synchronized larval staging to track the time required for lamellocyte differentiation to start in the lymph gland. During the late first-instar stage (18 hours <u>a</u>fter larval <u>h</u>atching, ALH), no lamellocytes were present (**Figures 1V, W, ZF,** with the GFP channel in **Figures S1C, D**). However, lamellocytes began to appear during the early second instar stage (24 hours ALH), without altering the size of the lymph gland (**Figures 1X, Y, ZF** and **Figures S1E, F**) till the late second instar stage (36 hours ALH) (**Figures 1Z, ZA, ZF**, and **Figures S1G, H**). By the early third instar stage (48 hours ALH), the lymph glands were beginning to disintegrate, resulting in smaller glands than the control lymph glands, which contained a large number of lamellocytes (**Figures 1ZB, ZC, ZF, and S1I, J**). Lamellocytes had begun to differentiate, with their numbers increasing rapidly during the wandering third larval stage. By the mid third-instar stage (60 hours ALH), Mys staining remained restricted to the medullary zone progenitors in controls, whereas *Gadd45^OE^* lymph glands showed strong ectopic Mys-expressing lamellocyte in the cortical zone and marked lobe disintegration (**Figures 1ZD, ZE, ZF,** and **Figures S1K, L**). These data indicate that Gadd45 overexpression causes precocious differentiation during lymph gland development.

We next examined published single-cell transcriptomic datasets (Cattenoz et al., 2020; Cho et al., 2020; Girard et al., 2021; Tattikota et al., 2020), which indicate that *Gadd45* transcripts are present at low levels across multiple blood-cell populations (**Figure S1M**). In third-instar lymph glands, RNA *in situ* hybridization detected endogenous *Gadd45* transcripts, with a stronger signal in Lozenge-positive (Lz⁺) crystal cells than in other cells (**Figures S1N-N’**). The absence of detectable signal in *Gadd45^F17^* mutant glands under the same conditions supports the specificity of the probe (**Figure S1O**). Additionally, a Gadd45 enhancer-trap Gal4 line (*Gadd45^149^*) driving GFP marked a subset of cells in the third-instar lymph gland (**Figures S1P–P**′). Together, these expression data suggest that Gadd45 is expressed in lymph gland blood cells and that its overexpression caused lamellocyte differentiation.

### Altered Gadd45 expression in the lymph gland core progenitor does not impact plasmatocyte differentiation

We next used the *Tep4-Gal4* driver to manipulate Gadd45 expression specifically in core progenitors (Girard et al., 2021; Goins et al., 2024) to determine whether this subset is sufficient to drive abnormal differentiation. Neither depletion nor overexpression of Gadd45 using the *Tep4-Gal4* driver recapitulated the phenotypes (**Figures 2A-D** and **Figures S2A-J**) that were observed in *dome^MESO^-Gal4,* such as lamellocytes differentiation and premature dispersal of primary lobes. Of note, *Tep4-Gal4*-driven *Gadd45*-expressing lymph gland showed a significant number of crystal cells (**Figures S2E, G, H**), which needs further experiments to find the reasons underlying the increased crystal cell phenotype, potentially activating Notch signaling for crystal cell differentiation (Duvic et al., 2002; Mukherjee et al., 2011).

**Figure 2:**
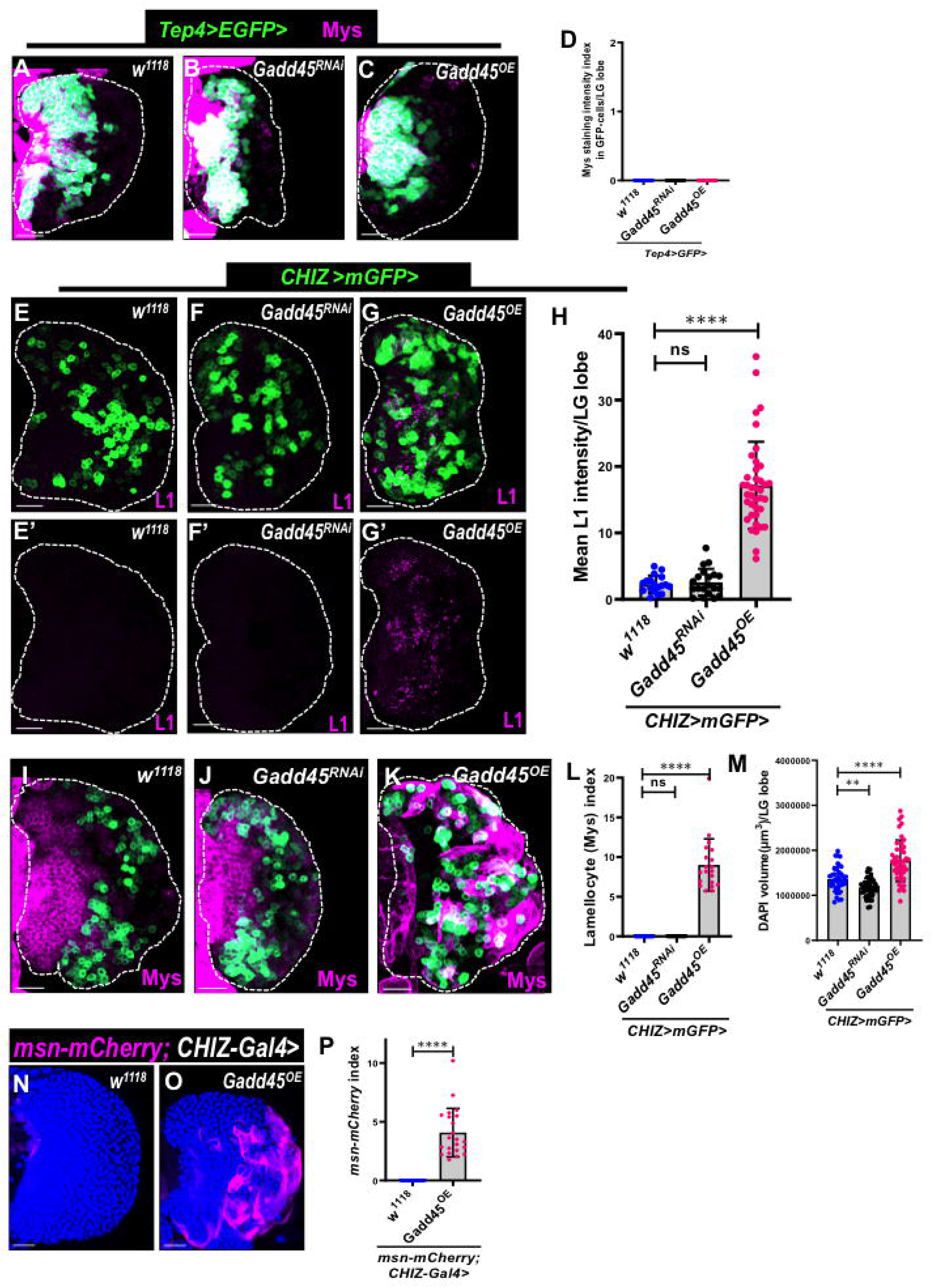
Overexpression of Gadd45 in lymph gland intermediate progenitors induces lamellocyte formation. **(A-C)** Core progenitor-specific *Tep4-Gal4, UAS-2XEGFP* (green) driven *Gadd45^RNAi^* (B, n=66) and *Gadd4^OE^* (C, n=69) do not induce any lamellocyte differentiation and look similar to wild-type (A, n=52) lymph gland. **(D)** Quantification of lamellocyte index in (A-C). **(E-G’)** Knockdown of Gadd45 in lymph gland intermediate progenitors (green) using *CHIZ-Gal4, UAS-mCD8::GFP* driver by expressing *Gadd45^RNAi^* (F-F’, n=20) shows no L1 (lamellocyte marker) immunofluorescence. However, *CHIZ-Gal4, UASmCD8::GFP* driven *UAS-Gadd45^OE^* (G-G’, n=38) shows lamellocyte differentiation marked by high L1 (magenta) compared to control (E-E’, n=18). **(H)** Mean L1 (lamellocyte marker) fluorescence intensity per anterior lymph gland lobe in (E-G’). **(I-K)** Intermediate progenitors (green) specific *CHIZ-Gal4, UAS-mGFP*-driven *UAS-Gadd45^RNAi^* (J, n=12) does not induce lamellocytes differentiation, while *UAS-Gadd45^OE^* (K, n=20) shows massive lamellocytes differentiation marked by Mys (magenta) compared to *w^1118^*(I, n=16). **(L)** Quantification of lamellocyte index in (I-K). **(M)** Quantification of lymph gland size using DAPI volume in (I-K). **(N-O)** *CHIZ-Gal4-*driven *UAS-Gadd45^OE^* (O, n=22) lymph glands show a vast number of *msn-mCherry*-positive (lamellocyte reporter, magenta) cells compared to control (*CHIZ-Gal4/+*) (N, n=16). **(P)** Quantification of lamellocyte index in (N-O). All confocal images represent maximum-intensity projections of the middle third optical sections of the wandering third instar lymph gland lobe from at least three independent biological experiments. White dashed lines outline a lymph gland lobe. Scale bar represents 25 μm. DAPI (blue)-stained nuclei. Error bars in graphs: mean ± S.D. ns.: not significant (p>0.01); **p<0.001; ****p<0.00001. n represents lymph gland lobe numbers.

### Gadd45 overexpression in intermediate progenitors of the lymph gland leads to lamellocytes differentiation

Next, we used the *CHIZ-Gal4* driver to manipulate Gadd45 expression only in intermediate progenitors (also called distal progenitors) (Spratford et al., 2021). As shown above, either depletion or overexpression of Gadd45 in core progenitors using *Tep4-Gal4* did not induce lamellocyte differentiation, nor did it alter overall lymph gland size or lobe integrity. Similarly, Gadd45 depletion in intermediate progenitors (*CHIZ>mGFP>Gadd45^RNAi^*) also failed to induce lamellocyte differentiation. (**Figures 2E-F’, H, I, J, L, M,** and **Figures S2K, L, N, O, P, R, S**). However, Gadd45 overexpression in intermediate progenitors (*CHIZ>mGFP>Gadd45^OE^*) resulted in a large number of lamellocytes differentiation, marked by L1 (Varga et al., 2019), msn-mCherry (Tokusumi et al., 2009), and Mys (Madhwal et al., 2020; Small et al., 2014) in the lymph gland (**Figures 2E-E’, G-G’, H, I, K, L, N-P**). However, no significant change was observed in Pxn+ cells (early plasmatocyte marker) and the number of crystal cells upon Gadd45 overexpression in intermediate progenitors (**Figures S2K, M, N, O, Q, R, S**). These results suggest that Gadd45 overexpression promotes lamellocytes differentiation, particularly in intermediate progenitors, thereby regulating the proper differentiation of mature blood cells. The unexpected appearance of lamellocytes in Gadd45 overexpression without an immune challenge indicates disrupted blood cell production. Therefore, in this study, we focus on how Gadd45 overexpression leads to a large number of lamellocytes.

### Gadd45 overexpression in circulating hemocytes causes lamellocyte differentiation

Expression of the *Hml*^Δ^*-Gal4* driver begins in intermediate progenitors and increases in mature blood cells in the lymph gland (Mondal et al., 2011; Spratford et al., 2021). As differentiating cells boost Hml expression, mature plasmatocytes exhibit higher levels of *Hml*^Δ^*-Gal4* driver (*Hml*^Δ^*>GFP*) expression. In contrast, crystal cells show relatively low expression (Mukherjee et al., 2011). Upon Gadd45 overexpression using the *Hml*^Δ^*-Gal4* driver, at 24 hrs ALH, *Hml*^Δ^*>Gadd45^OE^* lymph glands remain intact and comparable in size to control lymph glands, with no signs of enlargement or disintegration (**Figures S3A-B**). At 36 hrs ALH, *Hml*^Δ^*>Gadd45^OE^*lymph glands are also intact but enlarged compared to control (**Figures S3C-D**). By 48 hrs ALH (early L3), the lymph glands are completely disintegrated, with only a few lymph gland cells remaining (**Figures S3E-F**); at the wandering third-instar stage, the *Hml*^Δ^*>Gadd45^OE^*lymph gland shows hemocyte dispersal into circulation. *Hml*^Δ^*-Gal4* driver is also expressed in larval circulating blood cells(Sinenko & Mathey-Prevot, 2004) (**Figure 3A**). Therefore, we bled third-instar larvae to assay circulating hemocyte numbers, Hml>GFP+ cell numbers, and circulating lamellocytes numbers in the Gadd45 overexpression background and compare them with the control (**Figures 3A-E**). Remarkably, a high number of lamellocytes was observed in the circulation (**Figures 3A”-B”, E**), with a significantly increase in the total number of hemocytes and Hml>GFP+ cells index in the Gadd45-overexpressing background (**Figures 3A-D**). We also performed TUNEL staining to test whether cell death in circulating cells. Gadd45 overexpression increased total hemocyte number but reduced the TUNEL-positive cell index, with a slight decrease in *GFP*-positive cell proportion, further supporting that Gadd45 overexpression does not cause apoptosis (**Figures S3G-K**). These data suggest that Gadd45 overexpression in circulating hemocytes also causes lamellocytes differentiation.

**Figure 3:**
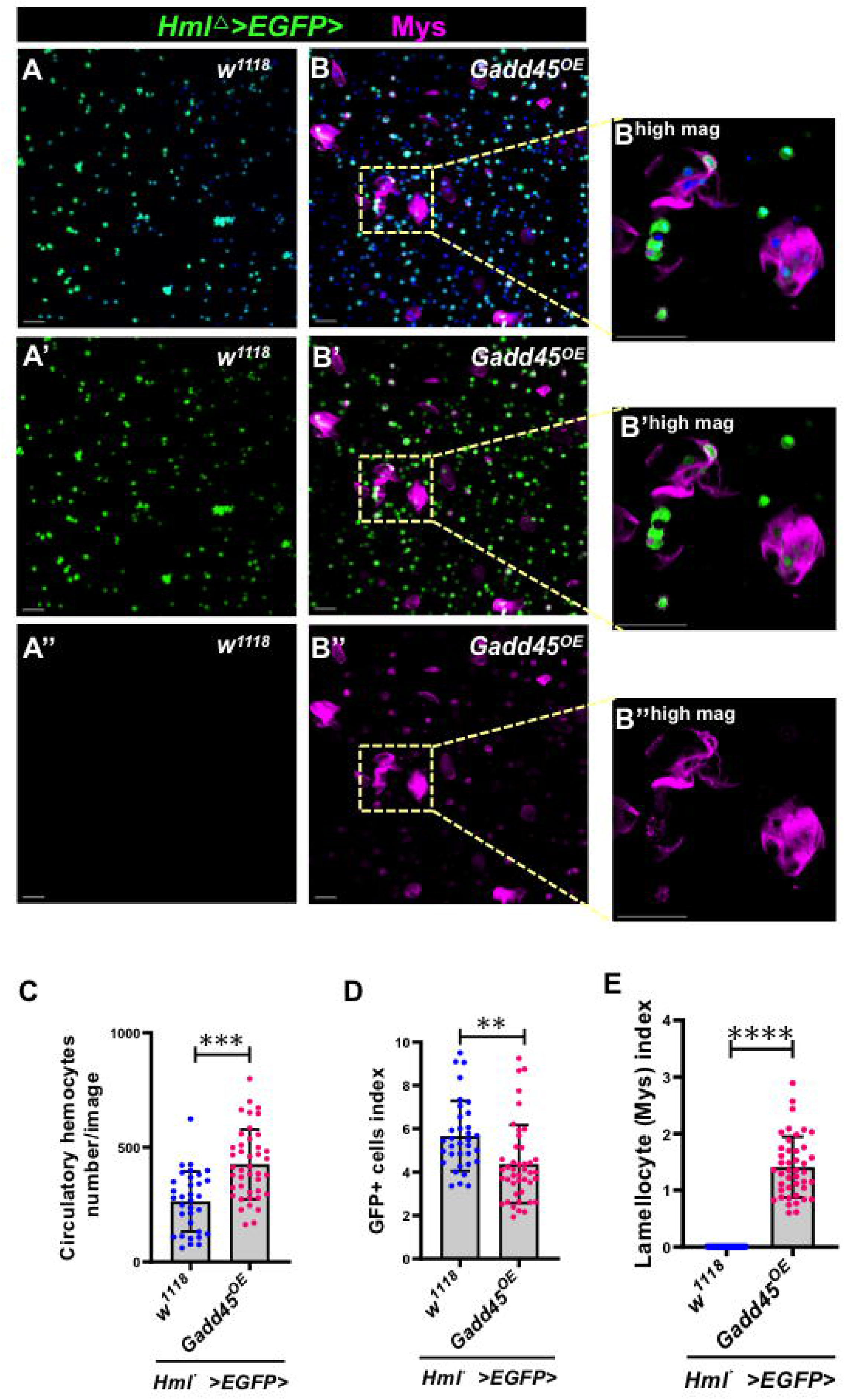
Gadd45 overexpression in circulating cells promotes lamellocyte differentiation. **(A-B’’)** *Hml*^Δ^*-Gal4, UAS-2xEGFP/+; Gadd45OE* (B-B’’, n=44) shows more Mys (magenta)-positive cells that mark lamellocytes among circulating cells of third-instar larvae than the control (*Hml*^Δ^*-Gal4, UAS-2xEGFP/+*) (A-A’’, n=36). **(C)** Quantification of circulating hemocytes per larva in (A-B’’). **(D)** Quantification of circulating GFP positive cell index in (A-B’’). **(E)** Quantification of circulating lamellocytes index in (A-B’’). All confocal images represent maximum-intensity projections of wandering third-instar circulating hemocytes from at least three independent biological experiments. Scale bar represents 50 μm. DAPI (blue)-stained nuclei. Error bars in graphs: mean ± S.D. ns.: not significant (p>0.01); **p<0.001; ****p<0.00001. n represents the number of larvae.

### Gadd45 overexpression activates the JNK, Toll, and JAK/STAT signaling pathways, causing lamellocyte differentiation

The intermediate progenitor zone in the third instar lymph gland expressed Jun Kinase (JNK) signaling pathway reporters, including extracellular matrix metalloprotease 1 (MMP1) and *puc-lacZ* (Maurya et al., 2024; Owusu-Ansah & Banerjee, 2009). Studies have also shown that lamellocyte differentiation requires activation of JNK signaling (Evans et al., 2022; Owusu-Ansah & Banerjee, 2009; Tokusumi et al., 2009). Interestingly, overexpression of Gadd45 in mammals and *Drosophila*’s various tissues is known to activate JNK signaling (Camilleri-Robles et al., 2019; Lu et al., 2001; Peretz et al., 2007). Therefore, we investigated whether Gadd45 overexpression in intermediate progenitors led to differentiation of lamellocytes due to the high activity of the JNK signaling pathway. Here, we used the *puc-lacZ* reporter and MMP1 immunostaining to assess JNK activity (Maurya et al., 2024; Owusu-Ansah & Banerjee, 2009; Uhlirova & Bohmann, 2006). Interestingly, Gadd45 overexpression in the intermediate progenitors (*CHIZ>mGFP>Gadd45^OE^*) resulted in a significant increase in *puc-lacZ*-positive cells (**Figures 4A-D**) and also a dramatic increase in MMP1 immunostaining in the third instar lymph gland compared to the controls and *Gadd45^RNAi^* backgrounds (**Figures S4A-D**). We were also able to partially rescue the lamellocytes phenotype of Gadd45 overexpression by co-expressing *bsk^DN^ and bsk^RNAi^* (*Drosophila JNK,* known as *basket*, *bsk*) (**Figures 4E-I** and **Figures S4E-I**). These results suggest that Gadd45 overexpression hyperactivated the JNK pathway, potentially triggering lamellocyte differentiation.

**Figure 4:**
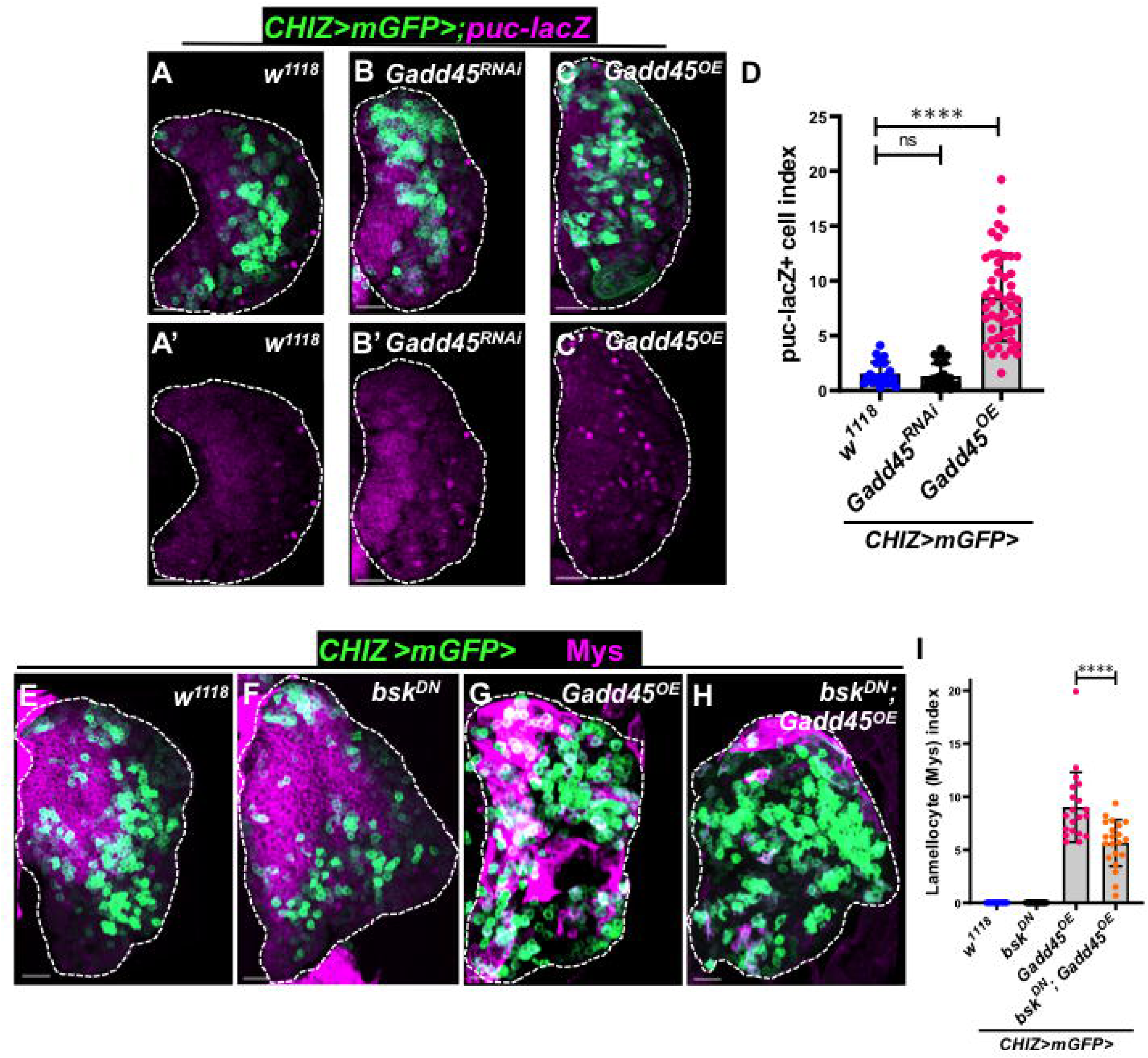
Gadd45 overexpression triggers JNK signaling activation in the lymph gland. **(A-C’)** The number of JNK signaling reporter *puc-lacZ* (magenta) positive cells remains unchanged in *CHIZ-Gal4, UAS-mCD8::GFP* driven *UAS-Gadd45^RNAi^* (B-B’, n=24) lymph gland lobes compared to control (*CHIZ>mGFP/+*) (A-A’, n=24), but expressing *UAS-Gadd45^OE^* (B-B’, n=49) shows a significantly increased number of *puc-lacZ*-positive cells. **(D)** Quantification of *puc-lacZ-*positive cell index in (A’-C’). **(E-H)** *CHIZ-Gal4, UAS-mCD8::GFP*-driven *bsk^DN^* (F, n=22) does not induce lamellocytes differentiation similar to the *w^1118^* (E, n=14), while expression of *UAS-Gadd45^OE^* (G, n=20) resulted in a large number of lamellocytes (magenta). Expression of *UAS-Gadd45^OE^* in the background of *bsk^DN^* in the intermediate progenitor reduces *Gadd45^OE^*-mediated lamellocytes (magenta) significantly (H, n=19). **(I)** Quantification of lamellocyte index in (E-H). All confocal images are maximum-intensity projections of the middle third of the optical sections of the wandering third-instar lymph gland lobe from at least three independent biological experiments. White dashed lines outline a lymph gland lobe. Scale bar represents 25 μm. Error bars in graphs: mean ± S.D. ns.: not significant (p>0.01); ****p<0.00001. n represents lymph gland lobe numbers.

After sterile injury to the larval cuticle, the inflammatory response activated the Toll/Dorsal pathway in the lymph gland without pathogenic infection and also caused lamellocytes differentiation (Evans et al., 2022). We ask whether this is also true in the context of Gadd45 overexpression in the lymph gland blood progenitors. The Toll signaling pathway’s downstream transcription factor, the Dorsal (*Drosophila* NF-κB) protein, is predominantly cytoplasmic in lymph gland progenitor cells (**Figures 5A-A’** and higher magnification **A-A’**) (Evans et al., 2022). Upon Gadd45 overexpression, Dorsal protein is predominantly nuclear in most third-instar lymph gland cells (**Figures 5B-B’** and higher magnification **B-B’**). Thus, Dorsal modulates the inflammatory response associated with Gadd45 overexpression in the larval lymph gland, possibly promoting lamellocyte differentiation.

**Figure 5.**
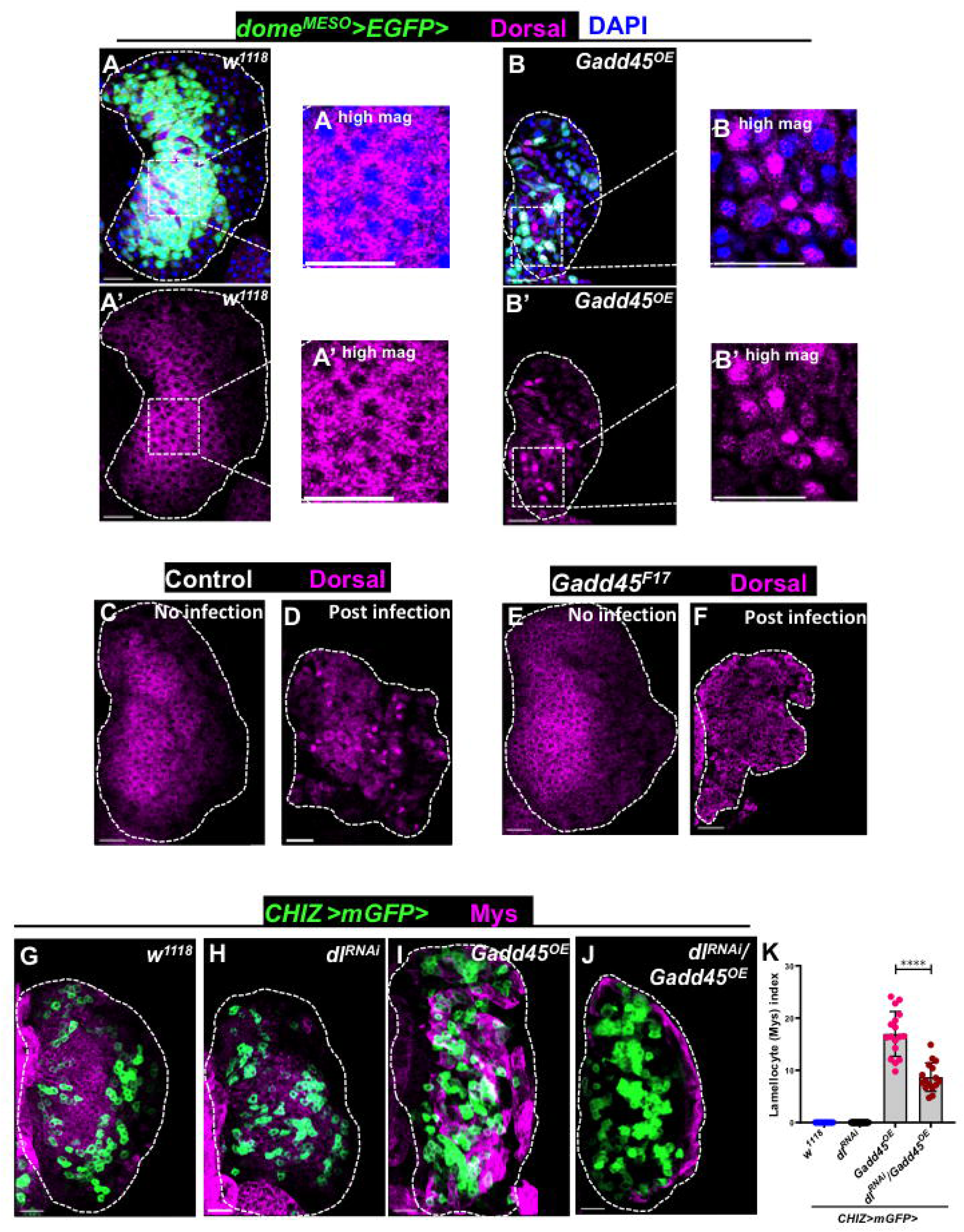
Gadd45 overexpression activates Toll signaling in the lymph gland. **(A-B’)** Compared with the control sets (*dome^MESO^-Gal4, UAS-2XEGFP/+*) (A-A’, n=22), the Toll signaling pathway was activated in *dome^MESO^-Gal4, UAS-2xEGFP* driven *UAS-Gadd45^OE^* (B-B’, n=54) expressing lymph glands, marked by nuclear Dorsal (magenta) staining. **(C-F)** Control lymph glands immunostained for Dorsal (magenta). Under uninfected conditions, Dorsal shows diffuse cytoplasmic expression (C, n=9). Post-wasp infection, Dorsal translocates to the nucleus (D, n=26), indicating Toll pathway activation. In *Gadd45^F17^* mutant lymph glands, the no-infection condition shows cytoplasmic Dorsal (E, n=14), similar to the control, whereas after wasp infection, Dorsal fails to translocate to the nucleus and remains cytoplasmic (F, n=44), indicating impaired activation of Toll signalling in the absence of Gadd45. **(G-J)** Control *CHIZ>mGFP/+* (G, n=8) and *CHIZ-Gal4, UAS-mCD8::GFP>UAS-dl^RNAi^* (H, n=20) alone shows no lamellocyte differentiation whereas *CHIZ-Gal4, UAS-mCD8::GFP>UAS-Gadd45^OE^* (I, n=17) shows robust lamellocyte formation. Co-expression of *dl^RNAi^* with *Gadd45^OE^* (J, n=16) rescues *Gadd45^OE^*-induced lamellocyte phenotype. **(K)** Quantification of lamellocyte index showing significant rescue *in CHIZ>mGFP>dl^RNAi^; Gadd45^OE^* (J) compared to *CHIZ>mGFP>Gadd45^OE^*(I). Control *CHIZ>mGFP/+* (G) and *CHIZ>mGFP>dl^RNAi^*(H). Confocal images represent maximum-intensity projections of the middle third optical sections of the wandering third instar lymph gland lobe from at least three independent biological experiments. White dashed lines outline a lymph gland lobe. Scale bar represents 25 μm. Error bars in graphs: mean ± S.D. ns.: not significant (p>0.01); ****p<0.00001. n represents lymph gland lobe numbers.

To further examine the role of endogenous Gadd45 in Dorsal localization, we compared control and *Gadd45^F17^* mutant lymph glands following *Leptopilina boulardi* wasp infestation. While Dorsal nuclear accumulation was evident in control lymph glands following wasp infestation, this response appeared impaired in *Gadd45^F17^* mutants (**Figures 5C–F**), suggesting that endogenous Gadd45 contributes to Dorsal activation following immune challenge.

To determine if Toll signaling is required for Gadd45-induced lamellocyte differentiation, we co-expressed *dorsal^RNAi^*along with Gadd45 overexpression in intermediate progenitors. While *dorsal^RNAi^*alone did not induce lamellocyte differentiation compared to control, co-expression of *dorsal^RNAi^* significantly reduced the elevated Mys-positive lamellocyte index observed in *Gadd45^OE^* lymph glands, indicating a partial rescue of the lamellocyte phenotype (**Figures 5G-K**).

Activation of JAK/STAT signaling is required for lamellocyte differentiation in larvae (Evans et al., 2022; Morin-Poulard et al., 2013). Here, we monitored an in vivo reporter that responds to nuclear STAT (*10XSTAT-GFP*) (Bach et al., 2007) and the JAK/STAT target gene *myospheroid* (*mys*) (Evans et al., 2022; Issigonis et al., 2009) by overexpressing Gadd45 in the intermediate progenitor background (*CHIZ-Gal4, 10XSTAT-GFP; UAS-Gadd45^OE^*). Remarkably, upregulated Mys+ cells in the cortical zone of the lymph glands also showed high *10XSTAT-GFP* reporter expression (**Figures 6A-B’** and **C-C’’’**, show higher magnification of Mys+ lamellocytes with high STAT-GFP). To examine the contribution of JAK/STAT signaling to Gadd45-driven lamellocyte differentiation, we co-expressed a dominant-negative *STAT92E* construct (*UAS-STAT^DN^*) together with Gadd45 in intermediate progenitors. Expression of *STAT^DN^* alone did not trigger lamellocyte differentiation relative to controls, whereas its co-expression with Gadd45 markedly reduced the increased Mys-positive lamellocyte index seen in *Gadd45^OE^*lymph glands, demonstrating a strong rescue of the lamellocyte phenotype (**Figures 6D-H**). Thus, overexpression of Gadd45-mediated lamellocyte differentiation in the lymph gland also leads to highly activated JAK/STAT signaling.

**Figure 6.**
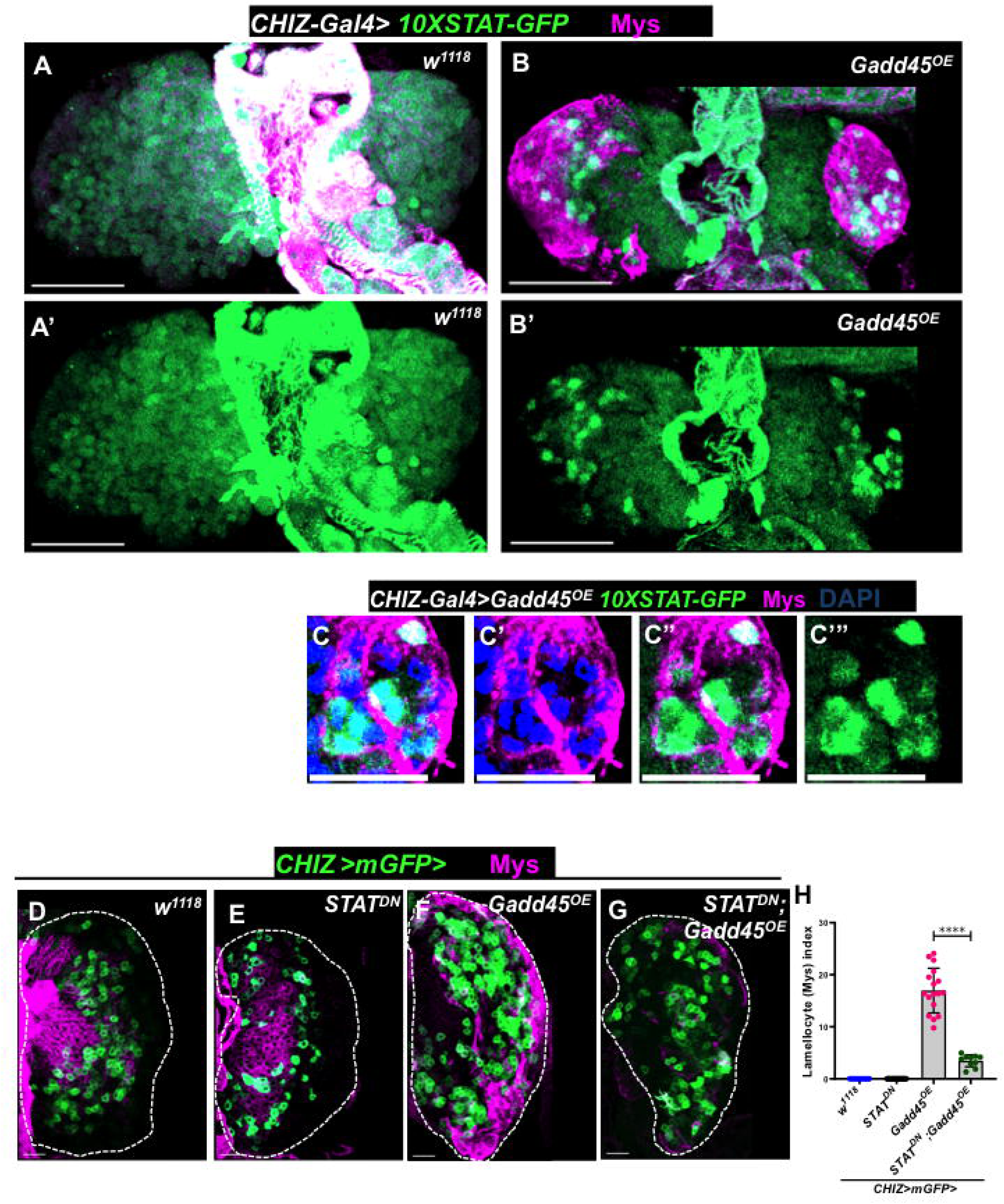
Gadd45 overexpression activates the JAK/STAT signaling pathway. **(A-B’)** *CHIZ-Gal4* driven *UAS-Gadd45^OE^* (B-B’, n=24) in *10X STAT-GFP* (STAT activity reporter) shows a high number of STAT-GFP positive cells in the differentiated zone of the lymph gland compared to control (*CHIZ-Gal4, UAS-mCD8::GFP/+*) (A-A’, n=12). **(C-C’’’)** Lymph gland lamellocytes in a Gadd45 overexpression background show a strong STAT-GFP (green) signal (High Magnification of B). **(D-G)** Control *CHIZ>mGFP/+* (D, n=8) shows progenitor-level Mys expression with no lamellocytes. Expression of *STAT^DN^* alone using the *CHIZ>mGFP* driver shows no ectopic lamellocytes (E, n=16). Overexpression of *Gadd45* using the *CHIZ>mGFP* driver induces massive lamellocyte differentiation with upregulated Mys (F, n=17). Co-expression of *STAT^DN^* with *Gadd45^OE^* suppresses *Gadd45^OE^*-induced lamellocyte differentiation (G, n=10). **(H)** Quantification of lamellocyte index showing significant rescue in *CHIZ>mGFP>STAT^DN^; Gadd45^OE^* (G) compared to *CHIZ>mGFP>Gadd45^OE^* (F). Control *CHIZ>mGFP/+* (D) and *CHIZ>mGFP>STAT^DN^* (E). Confocal images represent maximum-intensity projections of the middle third optical sections in A-B’ and D-G, and single optical sections in C-C”’ of the wandering third instar lymph gland lobe from at least three independent biological experiments. White dashed lines outline a lymph gland lobe. Scale bar represents 25 μm. Error bars in graphs: mean ± S.D. ns.: not significant (p>0.01); ****p<0.00001. n represents lymph gland lobes.

### Gadd45 is also associated with crystal cell differentiation and maintenance

Our results showed that Gadd45 depletion in the progenitor population causes a significant decrease in crystal cell number (**Figures 1L-M, O**). However, depletion of Gadd45 using the *CHIZ-Gal4* driver did not result in a significant change in crystal cell numbers in the lymph gland (**Figures S2O-P, R**); this may be because the *Gal4* driver is expressed only in intermediate progenitors. Additionally, we observed higher Gadd45 transcript expression in *Lz-GFP*-positive cells in the larval lymph gland (**Figures S1M-M’**). To determine whether the weaker crystal cell differentiation phenotype observed following Gadd45 depletion with *CHIZ-Gal4* was due to driver strength, we examined another intermediate progenitor Gal4 driver, *Nplp2-Gal4* (Cho et al., 2024; Rai et al., 2026), which has broader expression than *CHIZ-Gal4*. Notably, Gadd45 knockdown using *Nplp2-Gal4* significantly reduced the crystal cell index, whereas the plasmatocyte index remained statistically unchanged (**Figures 7A-E**). Reductions in crystal-cell-associated measurements were also observed following Gadd45 knockdown with other drivers, including *e33c-Gal4*, although the magnitude of these effects varied, whereas the plasmatocyte index remained unchanged (**Figures S5A-H**).

**Figure 7.**
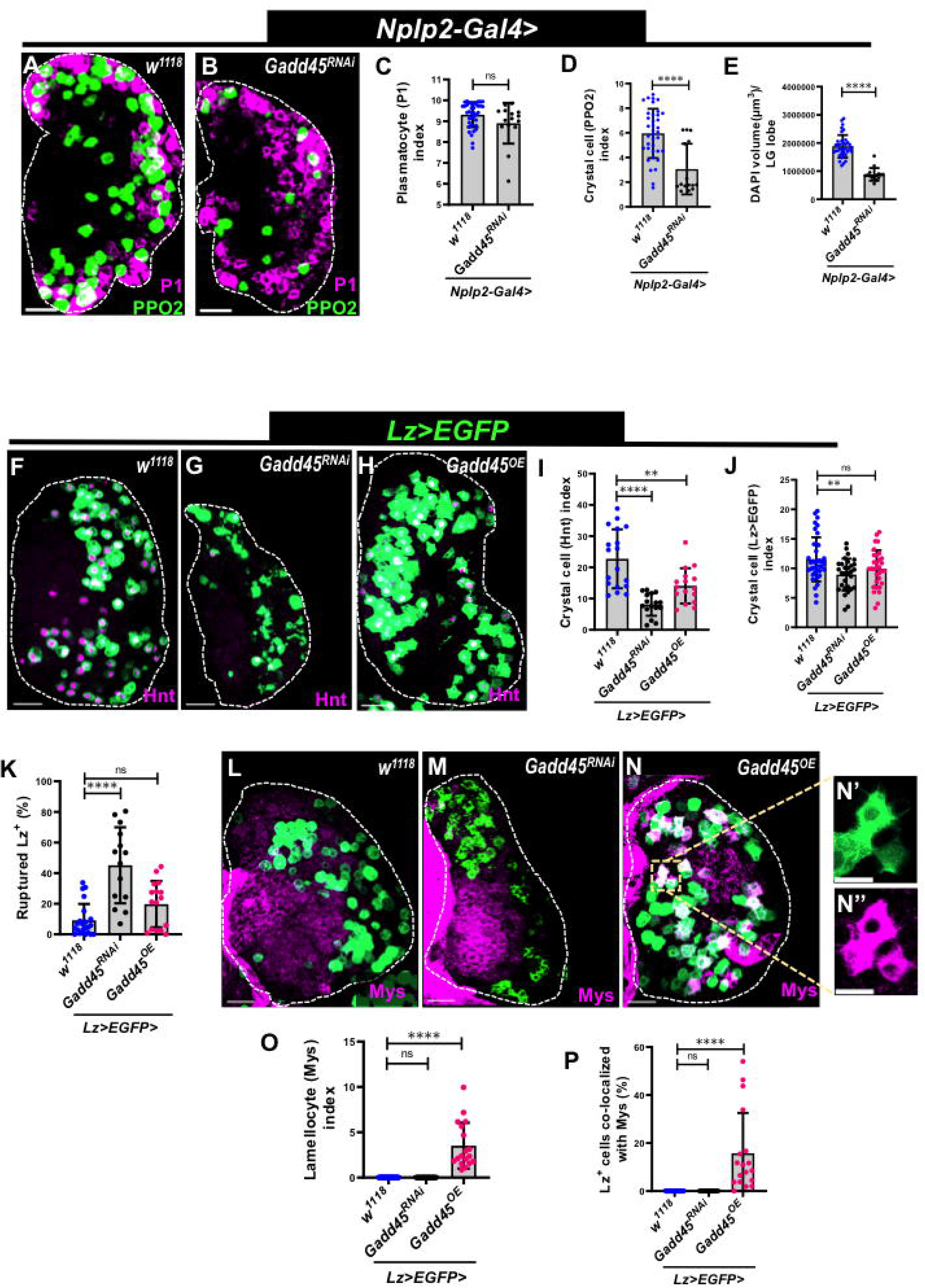
Gadd45 regulates crystal-cell-lineage-associated phenotypes and alters Mys expression in the lymph gland. **(A-B)** Knockdown of Gadd45 in lymph gland intermediate progenitors using the *Nplp2-Gal4* driver to express *Gadd45^RNAi^* (B, n=17), co-stained for P1+ plasmatocytes (magenta) and PPO2+ crystal cells (green), shows a strong reduction in crystal cells, while plasmatocytes remain comparable to control *Nplp2-Gal4/+* (A, n=37). **(C)** Quantification of plasmatocyte (P1) index in (A-B). **(D)** Quantification of crystal cell (PPO2) index in (A-B). **(E)** Quantification of lymph gland size using DAPI volume in (A-B). **(F-H)** Crystal cell-specific *Lz-Gal4, UAS-EGFP* (green) driven *Gadd45^RNAi^*(G, n=16) drastically decreases crystal cells marked by Hnt (magenta) and morphologically disrupted *Lz-GFP*-positive cells, whereas *Lz-Gal4, UAS-EGFP-*driven *Gadd45^OE^* (H, n=18) lymph glands show a slightly reduced Hnt index with no significant change in the *Lz+* cell number and morphology compared to control *Lz>EGFP/+* (F, n=18). **(I)** Quantification of crystal cell (Hnt) index in (F-H). **(J)** Quantification of crystal cell (*Lz>EGFP*) index in (F-H). **(K)** Quantification of morphologically disrupted *Lz-GFP*-positive cells (%) in (F-H). **(L-N)** Gadd45 was specifically knocked down (M, n=14) and overexpressed (N, n=18) in crystal cells using *Lz-Gal4, UAS-EGFP* (green). *Gadd45^RNAi^* shows no lamellocyte differentiation marked by Mys (magenta) and looks similar to control *Lz>EGFP/+* (L, n=22), whereas *Gadd45^OE^* shows ectopic lamellocyte differentiation. Inset in (N’-N”) shows high magnification of the boxed area with *Lz+* cells co-localized with Mys. **(O)** Quantification of lamellocyte (Mys) index in (L-N). **(P)** Quantification of *Lz+* cells co-localized with Mys (%) in (L-N). Confocal images represent maximum-intensity projections of the middle third optical sections except single optical sections in N-N” of the wandering third instar lymph gland lobe from at least three independent biological experiments. White dashed lines outline a lymph gland lobe. Scale bar represents 25 μm. Error bars in graphs: mean ± S.D. ns.: not significant (p>0.01); **p<0.001; ****p<0.00001. n represents lymph gland lobes.

We further tested whether Gadd45 deficiency or other genetic backgrounds affected crystal-cell abundance, as crystal-cell numbers also vary in different genetic backgrounds. We heat-shocked larvae at 60°C for 10 min, scored melanized (black-dot) crystal cells in third-instar larvae, and quantified black-dot structures per larva (Khadilkar et al., 2017) in *Hml*^Δ^*-Gal4/+*, *Gadd45^RNAi^/+*, *Gadd45^OE^/+*, and *Hml*^Δ^*>Gadd45^RNAi^*. Crystal-cell abundance was similar in all controls (**Figures S5I-M**). Together, these results suggest that Gadd45 depletion in the lymph gland affects crystal-cell-associated phenotypes depending on the cellular context in which its expression is altered.

We next examined the effects of altering Gadd45 expression in crystal cells using the *Lz-Gal4* driver to determine whether Gadd45 is also associated with crystal cell survival. Gadd45 knockdown reduced both the Hnt index (**Figures 7F-G, I**) and the Lz-positive cell index (**Figures 7F-G, J**), and was accompanied by an increased frequency of morphologically disrupted Lz-GFP-positive cells (**Figures 7F-G, K**). In contrast, Gadd45 overexpression using the *Lz-Gal4* driver resulted in a distinct set of phenotypes. The Lz-positive cell index remained relatively unchanged, whereas the Hnt index showed a modest reduction (**Figures 7F, H, I, J**). Notably, a subset of Lz-GFP-positive cells showed colocalization with the lamellocyte-associated markers Mys and L1 (**Figures 7L, M, N-N”, O, P** and **Figures S5N, P-P”**). These observations suggest that Gadd45 overexpression is associated with an altered cellular state, even in Lz-positive crystal cells.

Together, our findings show that Gadd45 is expressed in the Lz-positive population and that its depletion is associated with reduced crystal-cell-associated indices and changes in Lz-positive cell morphology. By comparison, Gadd45 overexpression is associated with lamellocyte differentiation. These results suggest that appropriate Gadd45 activity supports the normal development or maintenance of the crystal-cell lineage, whereas excessive expression alters the cellular state. Whether these changes reflect stress-associated lineage plasticity, altered differentiation, or another underlying mechanism remains to be investigated.

### Overexpression of Gadd45 alters the cell cycle status in the lymph gland

The Gadd45 family induces G2/M arrest by disrupting the cyclin/CDK complex (Vairapandi et al., 2002; Wang et al., 1999) and also binds to the cell cycle inhibitor p21, causing G1 arrest (Azam et al., 2001; Fan et al., 1999; Kearsey et al., 1995). Therefore, we examined whether Gadd45 overexpression in the lymph gland progenitors dysregulates the cell cycle status. At the third instar stage, seventy percent of the lymph gland progenitors are in the G2 phase (Cho et al., 2024; Goins et al., 2024; Kapoor et al., 2022), while only thirty-five percent of the intermediate progenitors (CHIZ+ cells) are in the G2 phase (Spratford et al., 2021). Overexpression of Gadd45 in progenitors (*dome^MESO^>FUCCI>Gadd45^OE^*) or intermediate progenitors (*CHIZ>FUCCI>Gadd45^OE^*) resulted in significant changes in cell cycle status (**Figures 8A, C, D,** and **Figures S6A-C**). Specifically, the number of G1-phase cells increased, while the number of G2-phase cells decreased in both zones (**Figures 8D** and **Figures S6C**). However, no significant change in cell cycle status upon Gadd45 depletion in progenitors (*dome^MESO^>FUCCI>Gadd45^RNAi^*) compared to the controls (**Figures 8A-B, D).** We also quantified mitotic cells in the third instar lymph gland using phospho-Histone H3 (pH3) immunostaining. No significant change in pH3+ cell numbers was observed in the lymph gland upon Gadd45 depletion in progenitors (*dome^MESO^>EGFP>Gadd45^RNAi^*) and intermediate progenitors (*CHIZ>mGFP>Gadd45^RNAi^*) compared to the controls (**Figures 8E-F’, H and I-J’, L**). However, overexpression of Gadd45 in the lymph gland progenitors (*dome^MESO^>EGFP>Gadd45^OE^*) and intermediate progenitors (*CHIZ>mGFP>Gadd45^OE^*) resulted in a significant increase in pH3+ cells (**Figures 8E-E’, G-G’, H and I-I’, K-K’, L**), indicating that more cells are mitotically active in the Gadd45-overexpressing background.

**Figure 8.**
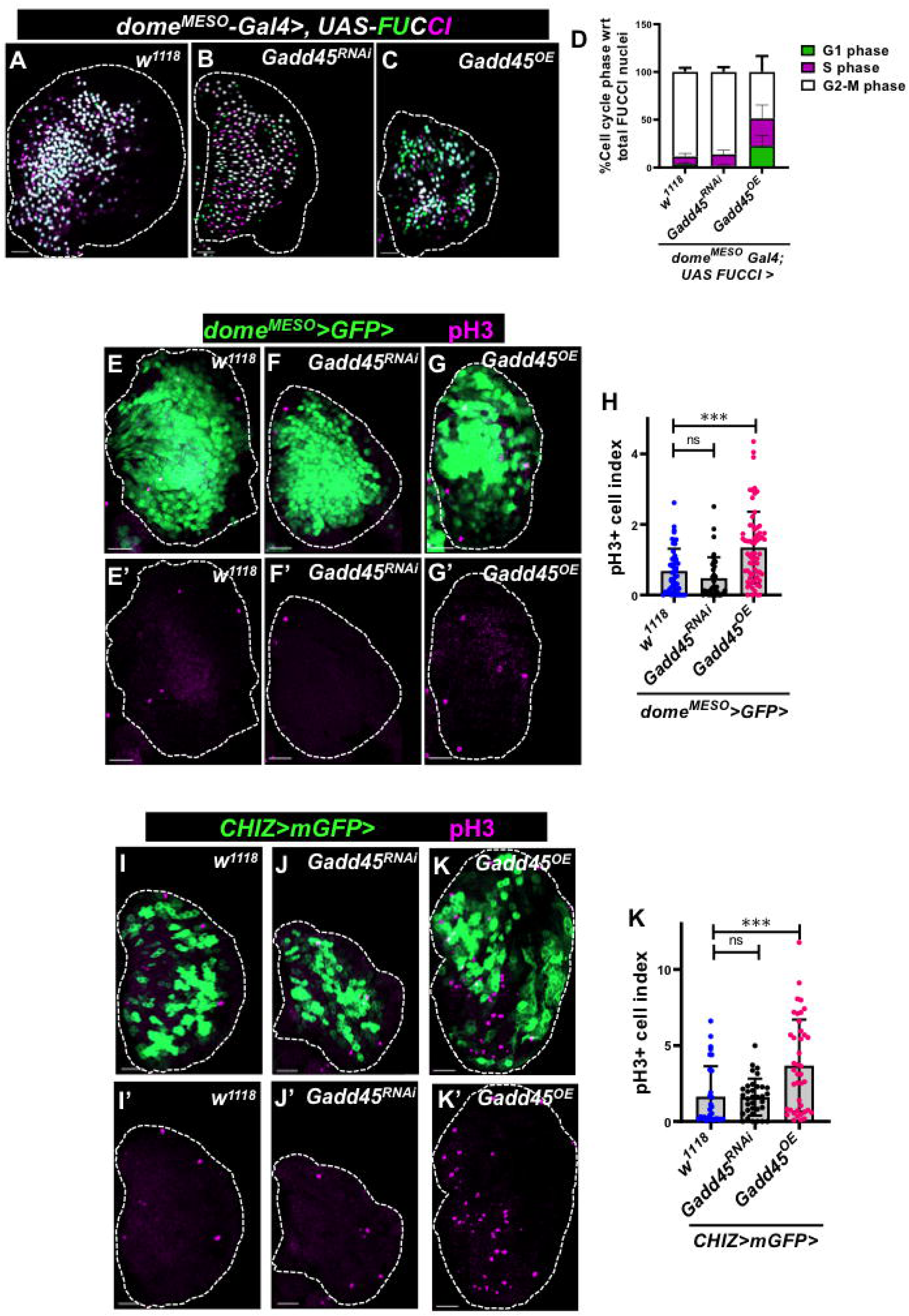
Cell cycle status of the lymph gland upon Gadd45 manipulation. **(A-C)** Cell cycle profile of progenitor cells tracked using Fly-FUCCI. Progenitor-specific expression of Gadd45 (*dome^MESO^-Gal4 UAS-FUCCI* driven *UAS-Gadd45^OE^)* (C, n=22) induces G1 phase arrest compared to knockdown of Gadd45 (*dome^MESO^-Gal4 UAS-FUCCI* driven *UAS-Gadd45^RNAi^)* (B, n=16) and control (*dome^MESO^-Gal4 UAS-FUCCI/+*) (A, n=13) (G1 phase: green, S-phase: magenta, G2/M phase: white). **(D)** Quantification of %G1, %S, and %G2/M cells in (A-C). **(E-G’)** Knockdown of Gadd45 in the intermediate progenitors (green) using *CHIZ-Gal4, UAS-mCD8::GFP* driven *UAS-Gadd45^RNAi^* (F-F’, n=34) has pH3-positive cells (a mitotic marker, in magenta) similar to the control (*CHIZ-Gal4, UAS-mCD8::GFP/+*) (E-E’, n=25). In contrast, *UAS-Gadd45^OE^*(G-G’, n=38) shows increased pH3-positive cells. **(H)** Quantification of pH3-positive cell index in (E-G’). **(I-K’)** Expression of Gadd45 in the progenitor cells using *dome^MESO^-Gal4, UAS-2xEGFP* driven *UAS-Gadd45^OE^* (K-K’, n=64) resulted in an increased number of pH3-positive cells (magenta), whereas knockdown of Gadd45 (J-J’, n=37) does not show any change in the pH3-positive cells compared to the control sets (E-E’, n=50). **(L)** Quantification of pH3-positive cell index in (I-K’). All confocal images represent maximum-intensity projections of the middle third optical sections of the wandering third instar lymph gland lobe from at least three independent biological experiments. White dashed lines outline a lymph gland lobe. Scale bar represents 25 μm. Error bars in graphs: mean ± S.D. ns.: not significant (p>0.01); ***p<0.0001. n represents lymph gland lobes.

Furthermore, we investigated whether lymph gland size is affected by apoptosis resulting from altered Gadd45 expression in lymph gland progenitors, as both knockdown and overexpression lead to smaller lymph glands, as shown above. TUNEL assay in Gadd45-depleted lymph gland progenitors showed similar numbers of TUNEL+ cells to those in the controls (**Figures S7A-B’, D**). However, the Gadd45 overexpression background showed fewer TUNEL+ cells than the control (**Figures S7A-A’, C-C’, D**), possibly due to the lymph gland dispersal phenotype. These findings suggest that altered Gadd45 expression in myeloid-type progenitor cells does not cause apoptosis, whereas Gadd45 overexpression alters their cell-cycle status.

Collectively, loss of the stress-response gene Gadd45 resulted in a smaller lymph gland, reduced crystal cell differentiation, and impaired crystal cell maintenance, whereas plasmatocyte differentiation remained unchanged. In contrast, Gadd45 overexpression altered cell-cycle status and induced lamellocyte differentiation by activating a combination of the Toll, JNK, and JAK/STAT signaling pathways (Figure 9).

**Figure 9:**
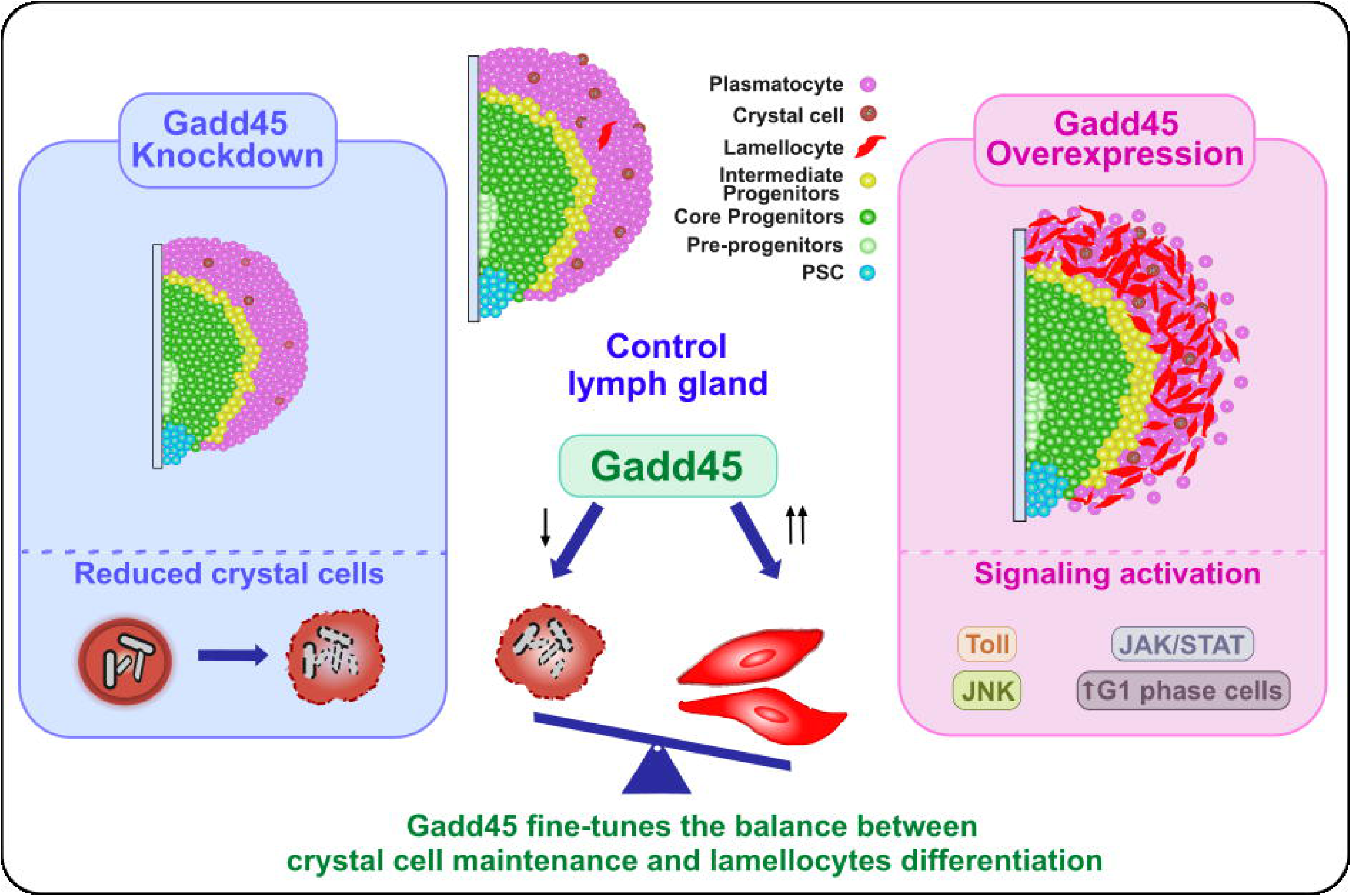
Schematic of Gadd45’s role in myeloid cell fate in the *Drosophila* larval lymph gland: Gadd45 knockdown reduces crystal cells, indicating it supports their maintenance. Gadd45 overexpression drives lamellocyte differentiation, disrupts gland organization, activates Toll, JNK, and JAK/STAT signaling, and induces G1 arrest. Thus, Gadd45 balances crystal cell maintenance and lamellocyte differentiation.

## Discussion

Studies reported altered Gadd45 expression in cancers like leukemia (Patel et al., 2022). Gadd45 is also induced by cytokines in myeloid differentiation, but its molecular mechanism remains unclear (Gupta et al., 2006). Earlier studies suggest that Gadd45 gene deficiency affects hematopoietic stem cells, warranting further study on myelopoiesis and hematopoiesis (Hoffman & Liebermann, 2007; Liebermann, 2022; Wingert et al., 2016).

Here, we report the role of Gadd45 in myeloid-type progenitor cells using the *Drosophila* lymph gland as an experimental model. We find that the absence of Gadd45 decreases the number of crystal cells, indicating that Gadd45 plays a role in crystal cell differentiation. The ruptured Lz⁺ cells observed after *Lz-Gal4*-driven *Gadd45^RNAi^* also suggest a possible role for Gadd45 in crystal-cell maintenance or survival. Crystal cells are rupture-prone, and Notch/HIF1α-associated mechanisms have been linked to their maintenance (Mukherjee et al., 2011). Mammalian studies have also linked HIF1α signaling to Gadd45α expression in cardiomyocytes (Krishnan et al., 2009), providing context for a possible connection between stress pathways and *Drosophila* Gadd45. Whether Gadd45 supports crystal-cell integrity in *Drosophila* through HIF1α (Sima) or a related mechanism remains to be investigated. Remarkably, plasmatocyte differentiation remains largely unaffected in Gadd45 loss, both in whole animals and in the blood system. Whether Gadd45-deficient differentiated plasmatocytes show similar phagocytic activity remains to be tested.

Functional redundancy among Gadd45 family members may mask phenotypes in single knockout mice. Although Gadd45α (Hollander et al., 1999; Wingert et al., 2016), Gadd45β (Gupta et al., 2005), and Gadd45γ (Hoffmeyer et al., 2001; Johnen et al., 2013) mutant mice were viable and exhibited no gross abnormalities in steady-state hematopoiesis, each knockout displayed distinct defects under conditions of genotoxic, inflammatory, or developmental stress. These observations suggest that Gadd45 proteins are largely dispensable for basal hematopoiesis but become critical during stress-induced cellular responses, with overlapping functions among family members contributing to functional compensation under homeostatic conditions. Gadd45α-Gadd45β double knockout mice exhibit partial embryonic lethality, neural tube defects, and impaired zygotic genome activation, highlighting their redundant roles in early embryonic development (Schule et al., 2019). Triple knockdown of Gadd45α/b/g in mouse embryonic stem cells produced a much stronger differentiation defect than single knockdowns, suggesting considerable redundancy among Gadd45 gene functions (Schule et al., 2019).

Gadd45 proteins are known stress sensors in mammals; Gadd45-deficient mice may be susceptible to immune compromise under various hematopoietic stress conditions (Gupta et al., 2005; Liebermann, 2022), including worse outcomes in chronic myeloid leukemia (Mukherjee et al., 2017; Sha et al., 2018). Although Gadd45-mediated myeloid-specific modulation of p38 and JNK signaling reduced the activity of granulocyte and macrophage chemotaxis in response to lipopolysaccharide (Salerno et al., 2012) this effect needs to be tested in *Drosophila* macrophages.

In *Drosophila*, Gadd45 overexpression has varying impacts, depending on the level and duration. Transient overexpression shows no apoptosis in the imaginal disc, whereas prolonged overexpression causes apoptosis and wing defects (Camilleri-Robles et al., 2019; Peretz et al., 2007). However, prolonged Gadd45 overexpression in the lymph gland myeloid-type progenitors does not kill them; instead, they differentiate into lamellocyte, suggesting that balanced Gadd45 is necessary for the appropriate number and types of differentiated blood cells in *Drosophila* larvae. Furthermore, *Drosophila* myeloid-type progenitors appear to be insensitive to stress response proteins like Gadd45. Third instar lymph gland progenitors are mainly arrested in the G2 phase of the cell cycle (Cho et al., 2024; Goins et al., 2024; Kapoor et al., 2022; Sharma et al., 2019). Gadd45 overexpression in blood progenitors alters the cell cycle status of third-instar lymph gland progenitors, with most progenitors entering the G1 phase rather than the G2 phase. This aligns with studies showing that Gadd45 directly interacts with cell cycle-related proteins and controls the cell cycle in the mammalian system (Vairapandi et al., 2002; Wang et al., 1999).

Our study shows Gadd45 overexpression causes the differentiation of lamellocytes - a response typically observed during the larval stage solely in reaction to immune challenges, injury, or wasp infestation. Activation of JAK/STAT signaling is also required for lamellocyte differentiation in larvae (Morin-Poulard et al., 2013). Upon larval injury, JAK/STAT signaling is also activated along with high expression of cytokine-like ligand unpaired-3 (Evans et al., 2022). Gadd45 overexpression in blood progenitors also activates the JAK/STAT signaling pathway, potentially leading to increased expression of unpaired-3, although this requires further investigation. Interestingly, Gadd45 overexpression initiates a differentiation program by activating the Toll/Dorsal and JNK signaling pathways. Thus, this study shows Gadd45 influences multiple signaling pathways in *Drosophila* myeloid-type cell development in the larval lymph gland, including the Toll, JNK, and JAK/STAT signaling pathways, which also play roles in innate immunity and injury-induced inflammation (Evans et al., 2022; Lemaitre & Hoffmann, 2007; Lemaitre et al., 1996; Stramer et al., 2008; Weavers et al., 2019).

Overall, Gadd45 overexpression in myeloid-type blood cells activates inflammatory responses through multiple signaling pathways in *Drosophila* larvae. Notably, a rare blood cell type emerges in the hematopoietic system, indicating that Gadd45 overexpression alone can induce lamellocyte formation without wasp parasitization. In the context of parasitization, injury, and mutant conditions, Toll- and JAK/STAT-dependent lamellocyte differentiation were reported (Evans et al., 2022; Harrison et al., 1995; La Marca & Richardson, 2020; Qiu et al., 1998). We show that this also applies to immune-stress-responsive Gadd45 overexpression. Our data extend earlier studies on Gadd45 regulating JNK signaling as a stress-responsive pathway across various contexts in both vertebrate and invertebrate model systems. Furthermore, this is the first report of a genetic association between Gadd45 and innate immune response signaling, such as the Toll and JAK/STAT pathways, in the *Drosophila* larval lymph gland.

Collectively, our findings indicate that Gadd45 supports normal crystal cell development and may promote an alternative lamellocyte-like state when overexpressed. Whether this effect reflects stress-related lineage plasticity, altered differentiation, or another mechanism remains unclear. Future research will clarify how *Drosophila* Gadd45 influences lamellocyte-specific gene expression and assess its relevance to the human myeloid system.

## Supporting information

Supplemental Figures with legends

## Acknowledgement

We thank Utpal Banerjee, Lolitika Mandal, Jiwon Shim, and Istvan Ando for providing reagents. We appreciate the Cytogenetics laboratory members and S.C. Lakhotia for their valuable input and support. We also thank the Bloomington Stock Center, USA, for fly stocks, and the Developmental Studies Hybridoma Bank, USA, for antibodies.

## Author contributions

Conceptualization, B.C.M.; methodology, B.C.M. and P.J.; investigation and analysis, P.J.; visualization, P.J. and B.C.M.; writing, B.C.M. and P.J.; funding acquisition and supervision, B.C.M.

During the manuscript preparation, we used Grammarly to check for errors in spelling, grammar, and clarity. The corresponding author then reviewed and edited the content and takes full responsibility for its accuracy.

## MATERIALS AND METHODS

### KEY RESOURCE TABLE

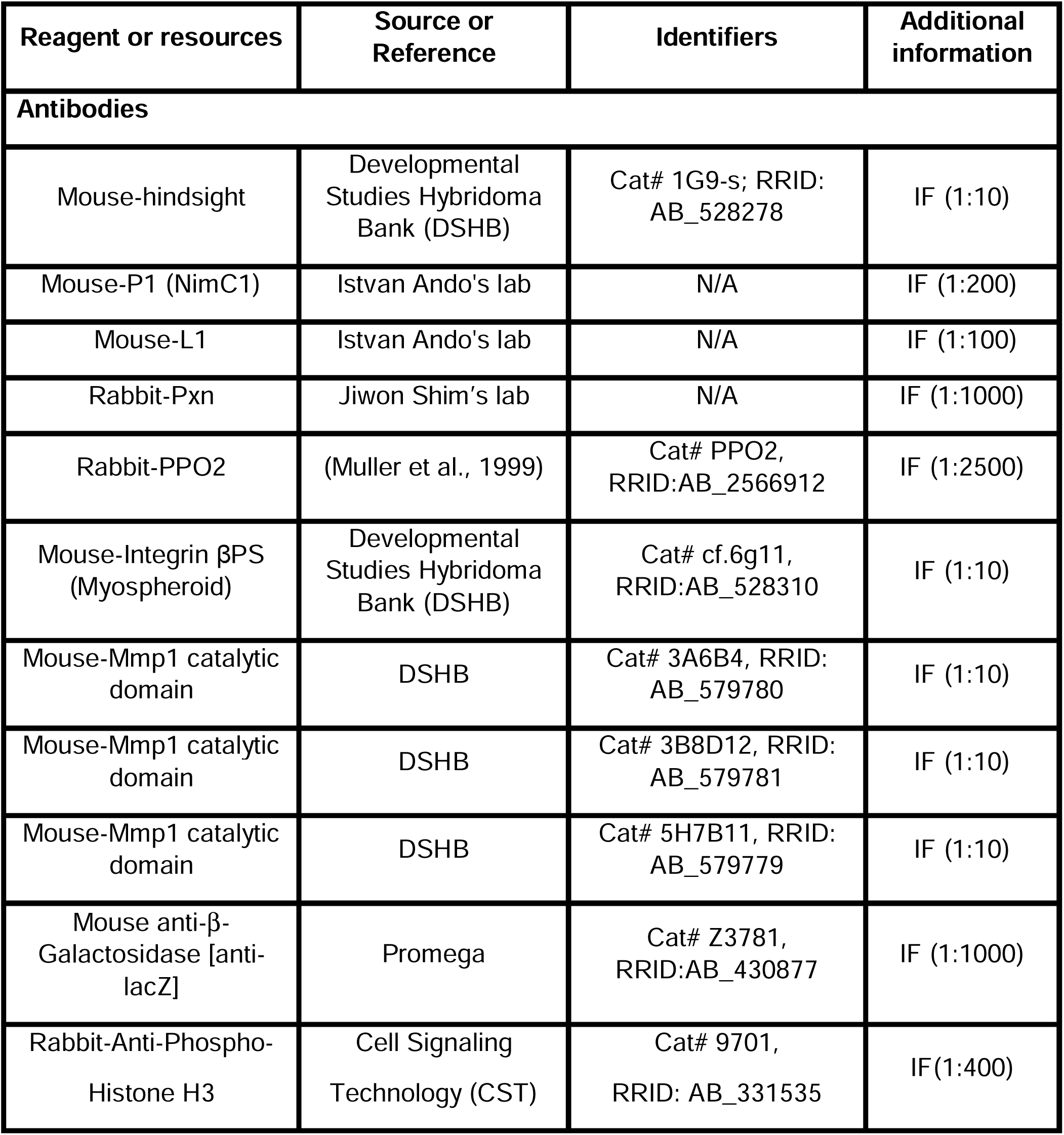
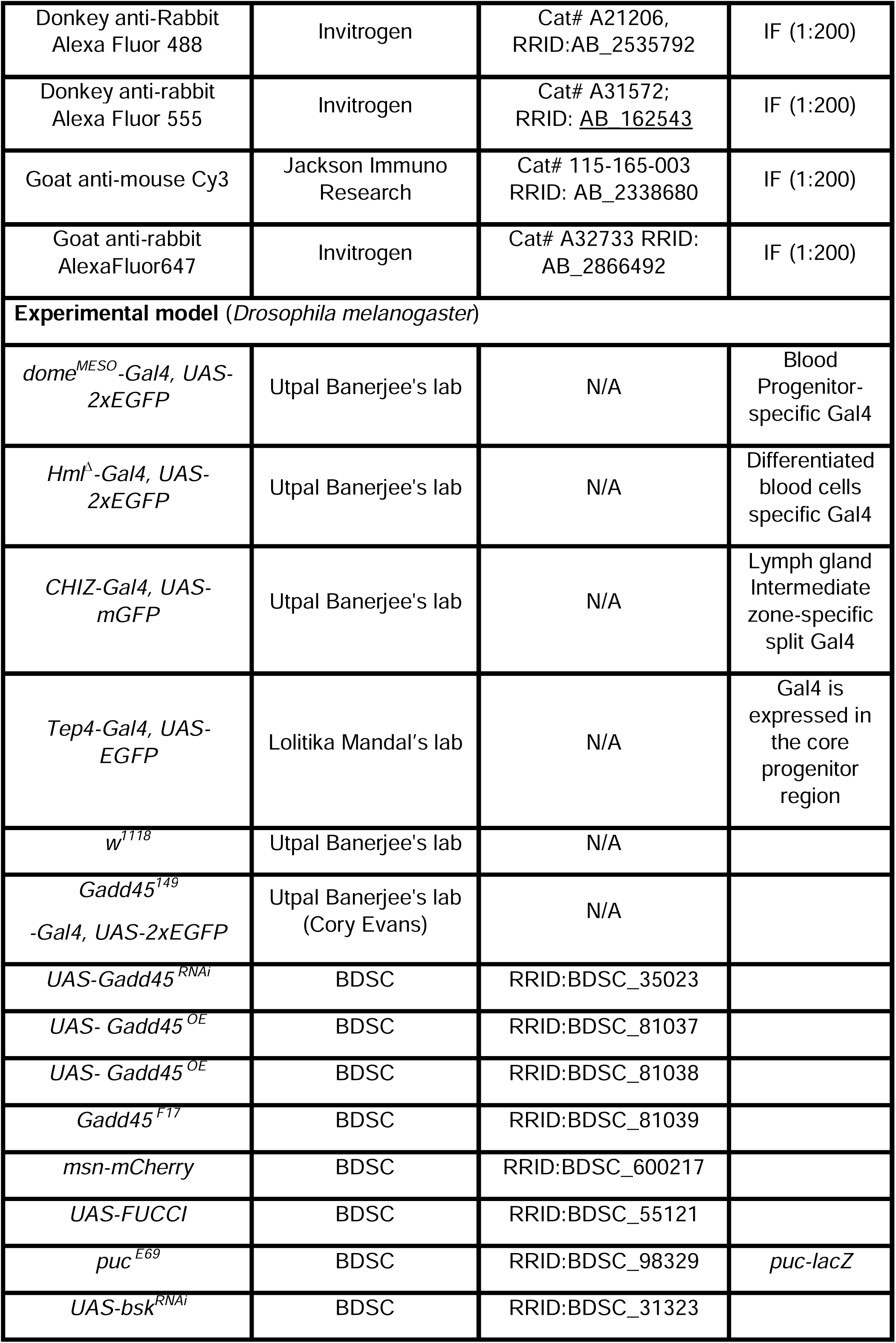
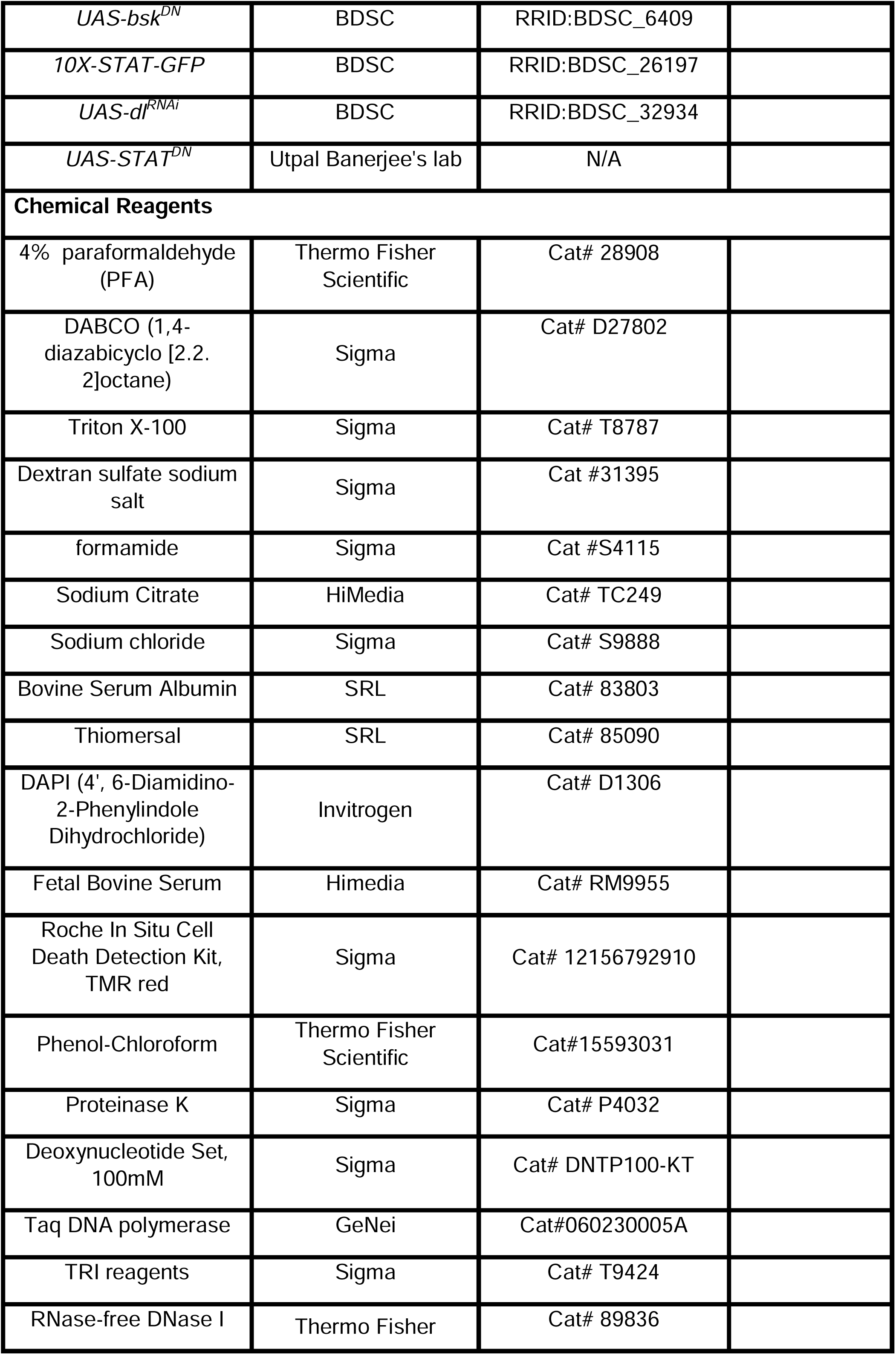
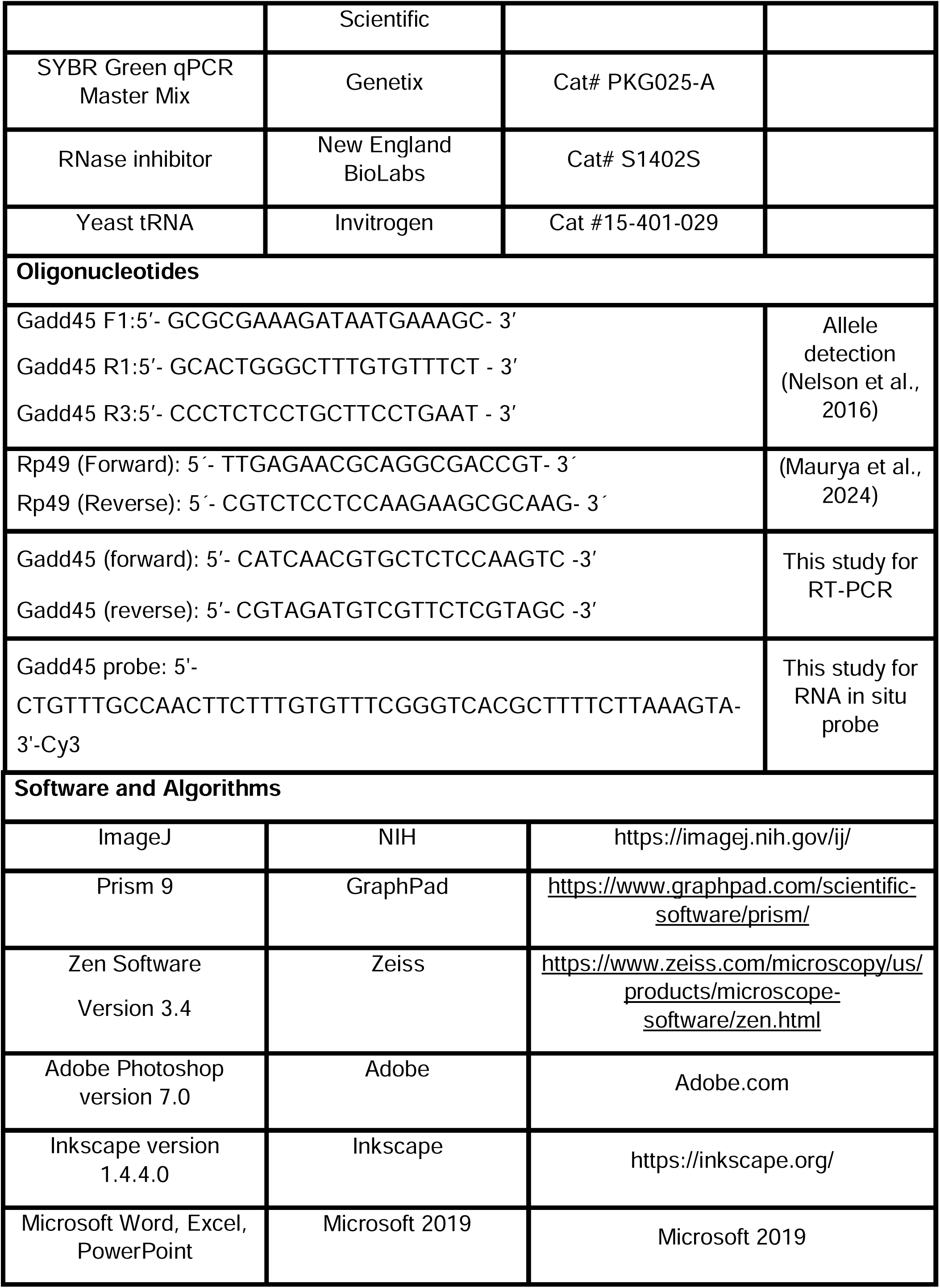

### EXPERIMENTAL MODEL AND STUDY PARTICIPANT DETAILS

*Drosophila* stocks were cultured using standard fly medium comprising 46 g/L cornmeal, 45 g/L sucrose, 18 g/L yeast extract, 7 g/L agar, supplemented with 3 mL/L propionic acid, and 3 g/L p-hydroxybenzoic acid methyl ester. In this study, these *Drosophila* stocks were used: *dome^MESO^-Gal4*, *dome^MESO^>2XEGFP*, *Hml*^Δ^*>2XEGFP*, *CHIZ>mGFP* (Spratford et al., 2021), *Gadd45^149^>EGFP, UAS-STAT^DN^* and *w^1118^*from Utpal Banerjee’s lab (UCLA). *Tep4>2XEGFP* from Lolitika Mandal (IISER Mohali). The following fly lines were obtained from Bloomington *Drosophila* Stock Center (BDSC): *UAS-Gadd45^RNAi^* (BL35023), *UAS-Gadd45^OE^*(BL81037, BL81038), *Gadd45^F17^* (BL81039), *msn-mCherry* (BL600217), *UAS-FUCCI* (BL55121), *puc^E69^* (BL98329), *UAS-bsk^RNAi^* (BL31323), *UAS-bsk^DN^* (BL6409), *UAS-dl^RNAi^* (BL32934).

All stocks and crosses were reared at 25°C, except those used in RNAi and Gal4-UAS-based expression experiments. The crosses involving RNAi lines were maintained at 29°C to enhance the efficacy of the Gal4-UAS RNAi system. Most experiments used lymph glands from wandering third-instar larvae; however, some used lymph glands from earlier stages at specified ages. Genetic crosses involved two-day-old virgin females and males post-eclosion. We used both sexes, and our study could not distinguish between them.

## METHODS DETAILS

### *Drosophila* larval lymph gland dissection and Immunostaining

Wandering third-instar larval lymph glands were mainly used for this study. The lymph glands were dissected in ice-cold 1xPBS (phosphate-buffered saline). After dissection, the tissues were fixed in 4% paraformaldehyde (PFA) (Thermo Fisher Scientific, Cat# 28908) for 30 minutes in microcentrifuge tubes. All steps were performed using a gentle-shaking rotator. After fixation, the tissues were washed thrice with 0.3% PBST (0.3% Triton X-100 in 1X PBS) for 10 minutes each, then kept in blocking solution (0.1% Triton X-100, 0.1% BSA, 10% Fetal Calf Serum, 0.1% sodium deoxycholate and 0.02% Thiomersal, in 1X PBS) for 30 minutes. Thereafter, samples were incubated at 4°C overnight with the primary antibody (diluted in blocking solution). The following day, the tissues were washed three times with 0.3% PBST, then reblocked with the blocking solution for 30 minutes. Then, the tissues were incubated in the secondary antibody for 2 hours at room temperature. Then, tissues were again washed thrice in 0.3% PBST, followed by counterstaining with DAPI (40,6-Diamidino-2-Phenylindole, Dihydrochloride, Thermo Fisher Scientific, Cat# D1306) (1 μg/mL). Finally, slides were mounted in DABCO (1,4-diazabicyclo [2.2.2] octane, Sigma, Cat# D27802, 2.5% DABCO in 70% glycerol made in 1X PBS)(Maurya et al., 2024).

These primary antibodies were used for immunohistochemistry: mouse anti-Hnt (1:10, 1G9-s, DSHB), mouse anti-P1 (1:200, I. Ando), mouse-L1 (1:100, I. Ando), rabbit-Pxn (1:1000, Jiwn Shim), rabbit anti-PPO2 (1:2500) (Muller et al., 1999), anti-Mys (1:10, CF.6G11, DSHB), anti-dorsal (1:10, anti-dorsal 7A4, DSHB), mouse anti-beta-Galactosidase [anti-lacZ] (1:1000, Z3781, Promega), mouse anti-MMP1 catalytic domain (a cocktail of three antibodies at dilution 1:10, 3A6B4, 3B8D12, 5H7B11, DSHB), and rabbit anti-phospho-Histone H3 (1:400, Cat# 9701, CST).

The secondary antibodies used for immunohistochemistry were goat anti-rabbit AlexaFluor647 (Invitrogen), donkey anti-rabbit Alexa Fluor 555 (Invitrogen), and goat anti-mouse Cy3 (Jackson Immuno Research). Secondary antibodies were used at 1:200 dilutions.

### Microscopy and image processing

All images were acquired using a Zeiss LSM900 confocal microscope with Zen Black software (version 3.4) under a 20x objective (digital zoom 1.0) and 40x/NA 1.3 objective (digital zoom 0.5x). A 2.0-micron optical section interval was used unless specified. The confocal imaging settings were kept identical for control and experimental samples. Each experiment was performed in biological triplicates with appropriate controls to ensure data reproducibility. Image processing used ImageJ software (NIH, USA) (available at ImageJ.nih.gov/ij). We used Adobe Photoshop (version 7) for the figure panel, and Inkscape (version 1.4.4.0) for the model preparation. To better represent lymph gland images, a maximum-intensity projection of the middle third of the lymph gland lobes was used to display the interior of the lymph gland. A white dotted line marks the boundary of the lymph glands.

### TUNEL staining for cell death assay

The lymph glands were isolated in cold 1X PBS, fixed in 4% paraformaldehyde for 30 minutes at room temperature, and washed thrice with PBST containing 0.3% Triton X-100. TUNEL (terminal deoxynucleotidyl transferase-mediated deoxyuridine triphosphate nick-end labeling) staining was performed using the In Situ Cell Death Detection Kit, TMR Red (Sigma, Cat# 12156792910), following the manufacturer’s instructions, as previously described (Maurya et al., 2024).

Circulating hemocyte TUNEL assay: Third-instar wandering larvae were bled into 20 μL PBS on a clean slide, and hemocytes were allowed to adhere for 30 minutes in a moist chamber. After removing the PBS, the cells were fixed with 4% PFA for 30 minutes. Following fixation, immunostaining and the TUNEL assay were performed as described above for the lymph gland.

#### *Drosophila* larval staging

Flies were allowed to lay eggs on agar plates for 12 hours to ensure synchronization. Afterward, the embryos were incubated at 25°C for another 12 hours. The hatched larvae were removed with a paintbrush, leaving the unhatched embryos to incubate at 25°C for 30 minutes (Mondal et al., 2011). Newly hatched larvae were carefully transferred to fresh food vials and incubated at 29°C. Larvae at 18, 24, 36, 48, and 60 hours after larval hatching (ALH) were collected for dissection and Mys staining. Lymph glands were isolated, and immunostaining was performed as specified in the immunostaining section.

#### *L. boulardi* parasitism assay

For wasp infestation assays, vials containing 40–50 larvae were exposed to four to five male and female *Leptopilina boulardi* wasps at 48 h after larval hatching (ALH). After 3–4 h of co-culture, wasps were removed, and *Drosophila* larvae were allowed to develop to the wandering third-instar stage before dissection (Madhwal et al., 2020). Infestation was confirmed by direct observation of wasp eggs in the hemolymph during dissection. All infestation procedures were carried out at 25°C.

### Heat-induced crystal cell melanization assay

Crystal cells were visualized by heat-induced melanization, following previously described methods(Frankenreiter et al., 2021; Khadilkar et al., 2017). Briefly, wandering third-instar larvae were collected, briefly washed in PBS, and transferred in batches of five into PCR tubes containing moist tissue paper. The tubes were subjected to heat treatment at 60°C for 10 min to induce crystal cell melanization. Following heat treatment, melanized crystal cells were identified as distinct black spots and imaged using a Zeiss stereomicroscope. The number of melanized crystal cells was quantified in the last two posterior abdominal segments of each larva using ImageJ.

### Quantification of Lymph gland phenotypes

All measurements were performed using ImageJ software (NIH, USA). The comprehensive approach for examining different parameters is outlined below:

### Lymph gland and zone volume analysis

To measure the volume of multichannel images, all image channels were first separated, and a single threshold was selected to best match the actual staining, ensuring consistency throughout the measurement. The thresholding method is used in image processing to select pixels of interest based on their intensity values. The lymph gland region was then marked using the freehand selection tool. Subsequently, the “Macro volume” plugin was used to determine the fluorescent area (for DAPI, GFP, Pxn, and P1) in each optical section. Subsequently, the fluorescent region in every optical section was summed and multiplied by the stack interval (2 μm) to calculate the volume.

The index for each marker in a lymph gland lobe was calculated as described below (Destalminil-Letourneau et al., 2021; Morin-Poulard et al., 2016):

For PPO2/Hnt/Mys/msn-mCherry/puc-lacZ/pH3/TUNEL, the index was calculated as (number of positive cells / total DAPI volume) × 100,000.

For progenitors, P1, and Pxn, which form regions where individual cells cannot be distinguished, we measured the total marker-positive volume. The index was calculated as (marker+ volume / total DAPI volume) × 10.

### L1 and MMP1 fluorescent intensity analysis

The mean fluorescent intensity of L1 and MMP1 staining was assessed as described in the literature (Kapoor et al., 2022; Louradour et al., 2017). For quantification, two middle z-stack sections from each lymph gland image were chosen. Signal intensity was measured within a fixed 50 × 50 μm square region using the “Measure” tool. For background correction, a 50 × 50 μm square outside the lymph gland was selected, and the mean background intensity was subtracted from the mean intensity of the lymph gland. For all intensity measurements, the laser setting for each specific experimental arrangement was maintained constant. Controls were examined simultaneously with the experiments each time.

### FUCCI area analysis

We used the Fly-FUCCI transgenic fly line (Zielke et al., 2014) for estimating the G1, S, and G2/M populations in the progenitor pool; the middle 2 optical sections that most accurately depicted the medullary zone of the obtained confocal image of the lymph gland at a specific developmental time point were merged (Shim et al., 2012). Next, we isolated the channels of the combined image to examine the green and magenta channels separately. To do this, FUCCI reporter-positive cells (green/magenta) were thresholded, then the “watershed” and “analyze particles” commands were applied to count green- and magenta-positive cells independently. To estimate the white population, the regions of interest for the green-positive cells were superimposed on the red channel image, and the "green only" cells were counted by hand. This “green only” value indicates the count of cells in the G1 phase that was subsequently subtracted from the overall green-positive cells measured to derive the yellow cells representing the G2/M population. The “magenta only” value, representing the S phase, was similarly obtained by removing yellow cells from the total calculated red cells. The values were subsequently normalized to the total number of FUCCI-positive cells from the merged stacks and expressed as percentages of the G1, S, or G2/M population (Kapoor et al., 2022).

### Circulating blood cell counting

Third instar wandering larvae from various genotypes were bled onto a clean slide using 20 μL of phosphate-buffered saline (PBS). Hemocytes were allowed to adhere to the glass slide for 30 minutes. Afterward, PBS was gently removed, and the cells were fixed with 4% PFA for 30 minutes. Following fixation, the hemocytes were washed twice with PBS, stained with DAPI, and then washed again with PBS. The samples were then mounted using DABCO mounting medium. Four non-overlapping fields of view were images were captured from each third-instar larval bleeding sample using a Zeiss confocal microscope with 20x objectives, through the 488 nm, 555 nm, and 405 nm channels. The numbers of DAPI-, Mys-, and GFP-positive hemocytes in each image were manually counted using the ImageJ plugin.

GFP and Mys indexes correspond to (Number of GFP/Mys marked cells/Total DAPI number per image) ×10.

TUNEL index corresponds to (TUNEL+ cells number/Total DAPI number per image) ×100.

### DNA extraction and PCR methods for the detection of Gadd45 deletion

To prepare genomic DNA, fifty flies are homogenized in a lysis buffer with Proteinase K to break down tissues and proteins. The mixture is then purified using the traditional phenol-chloroform extraction method. After purification, DNA is obtained via salt-ethanol precipitation. The DNA pellet is washed with ethanol to remove impurities, air-dried to eliminate residual solvent, then resuspended in water to produce concentrated, purified genomic DNA. This DNA is used as a template for PCR amplification of the *Gadd45* and *Rp49* genes. The PCR protocol included an initial denaturation at 95°C for 5 minutes, followed by 30 cycles of denaturation at 95°C and annealing at 56°C for 1 minute each, extension at 72°C for 1 minute 40 seconds, and a final extension at 72°C for 10 minutes. Rp49 served as an internal control, and PCR product sizes were analyzed by agarose gel electrophoresis. Primers for Gadd45 and Rp49 are listed below. Gadd45 F1:5’- GCGCGAAAGATAATGAAAGC- 3’ Gadd45 R1:5’-GCACTGGGCTTTGTGTTTCT - 3’ Gadd45 R3:5’- CCCTCTCCTGCTTCCTGAAT - 3’ Rp49 (Forward): 5’-TTGAGAACGCAGGCGACCGT- 3’ Rp49 (Reverse): 5’- CGTCTCCTCCAAGAAGCGCAAG-3’ To verify the *Gadd45^F17^* mutant created by Nelson et al. (2016), we used the same primer set: Gadd45_F1 / Gadd45_R1, to amplify the region deleted by the excised P-element. The *Gadd45^F17^* mutant line (BL81039) did not yield amplification with these primers; therefore, we tested it with the Gadd45_F1 / Gadd45_R3 primers to detect deletions. A smaller DNA fragment was observed with the Gadd45_F1 / Gadd45_R3 set compared to the control, indicating that the fly line *Gadd45^F17^* (BL81039) is homozygous mutated in the Gadd45 locus.

### Quantitative real-time PCR

Total RNA was extracted from seventy-eight wandering third-instar larval lymph glands using TRIzol reagent according to the manufacturer’s instructions (Sigma-Aldrich, Cat# T9424). The RNA pellets were resuspended in 12 μl of DEPC-treated Milli-Q water, and RNA concentration was determined by spectrophotometric analysis. Subsequently, 1 μg of total RNA from each sample was treated with 1 U of RNase-free DNase I (Thermo Fisher Scientific, Cat# 89836) at 37 °C for 30 min to remove any contaminating genomic DNA. First-strand cDNA was then synthesized from the DNase I-treated RNA samples following the standard cDNA synthesis protocol. Quantitative real-time PCR (qRT-PCR) was performed using 5 μl of SYBR Green qPCR Master Mix (Genetix, Cat# PKG025-A) and 2 pmol/μl of each primer in a final reaction volume of 10 μl on an ABI 7500 Real-Time PCR System. Relative changes in gene expression were determined using the comparative Ct (ΔΔCt) method, with Rp49 serving as the internal reference gene for data normalization. The primers used in this study are listed below: Gadd45 (forward): 5’-CATCAACGTGCTCTCCAAGTC -3’ Gadd45 (reverse): 5’-CGTAGATGTCGTTCTCGTAGC -3’ Rp49 (forward) 5’-TTGAGAACGCAGGCGACCGT-3’ Rp49 (reverse) 5’-CGTCTCCTCCAAGAAGCGCAAG-3’.

### RNA fluorescence *in situ* hybridization (RNA FISH)

A 47 nt Cy3-labeled antisense DNA oligonucleotide complementary to *Gadd45*: 5’-CTGTTTGCCAACTTCTTTGTGTTTCGGGTCACGCTTTTCTTAAAGTA-3’-Cy3 was synthesized commercially and purified by RNase-free HPLC (GenScript). Wandering third-instar larvae were dissected in chilled 1X PBS, fixed in 4% paraformaldehyde for 30 min at room temperature, and washed three times in 0.3% PBST. Tissues were permeabilized in 1% Triton X-100 with 1 U/µL RNase inhibitor (New England BioLabs, Cat # S1402S) for 20 min at 40°C, washed in 0.3% PBST and 2X SSC, and hybridized overnight at 50°C in buffer containing 50% formamide (Sigma, Cat #S4115), 2X SSC pH 7.0, 10% dextran sulfate (Sigma, Cat #31395), 1 mg/mL yeast tRNA (Invitrogen, Cat #15-401-029), and 5 ng/µL Cy3-labeled Gadd45 oligo probe. Post-hybridization washes were performed at 42°C in 50% formamide/2X SSC, 2X SSC, and 1X SSC, followed by 1X PBS washes at room temperature. Nuclei were counterstained with DAPI for 10 min; tissues were mounted in DABCO and imaged on a confocal microscope using appropriate filters.

### Statistical analysis

At least three independent biological replicates, each with three experimental repeats, were analyzed, and a representative image is shown in the figures. The quantifications include all datasets analyzed. Statistical tests were conducted using GraphPad Prism 9 and Microsoft Excel 2019. A two-tailed, unpaired Student’s t-test compared two groups, while a two-way ANOVA with Tukey’s post hoc test was used for multiple comparisons. All P-values presented represent statistical significance (**** if p<0.0001, *** if p<0.001, ** if p<0.01, and * if p<0.05), and ns for not significant p > 0.05. In the supplementary file, we have provided details of the statistical analysis for each set of experimental data.

