## Supplemental Figures with legends for "Gadd45 regulates fate decisions of myeloid-type blood cells in *Drosophila*"

### Figure S1

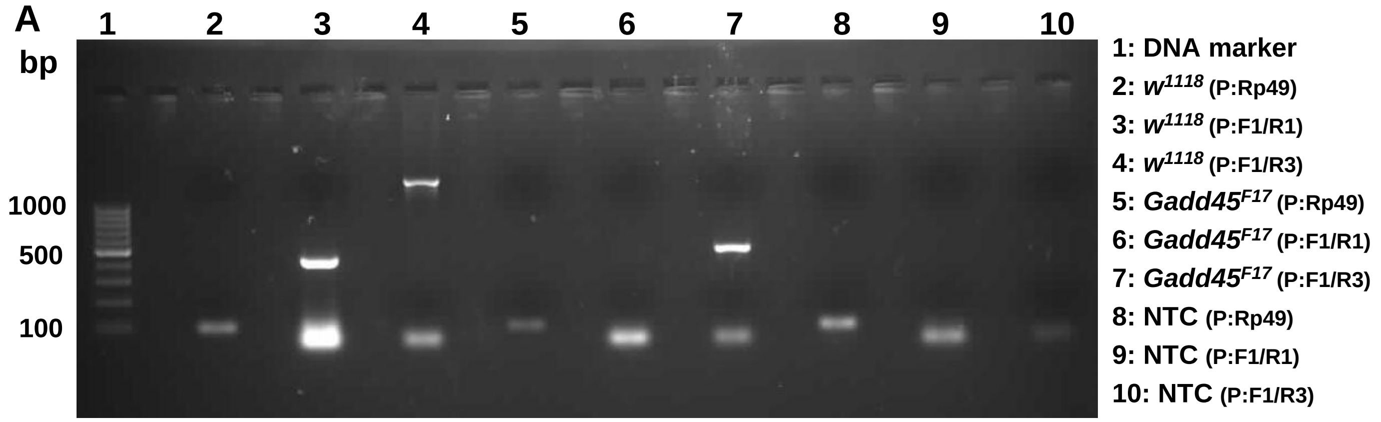


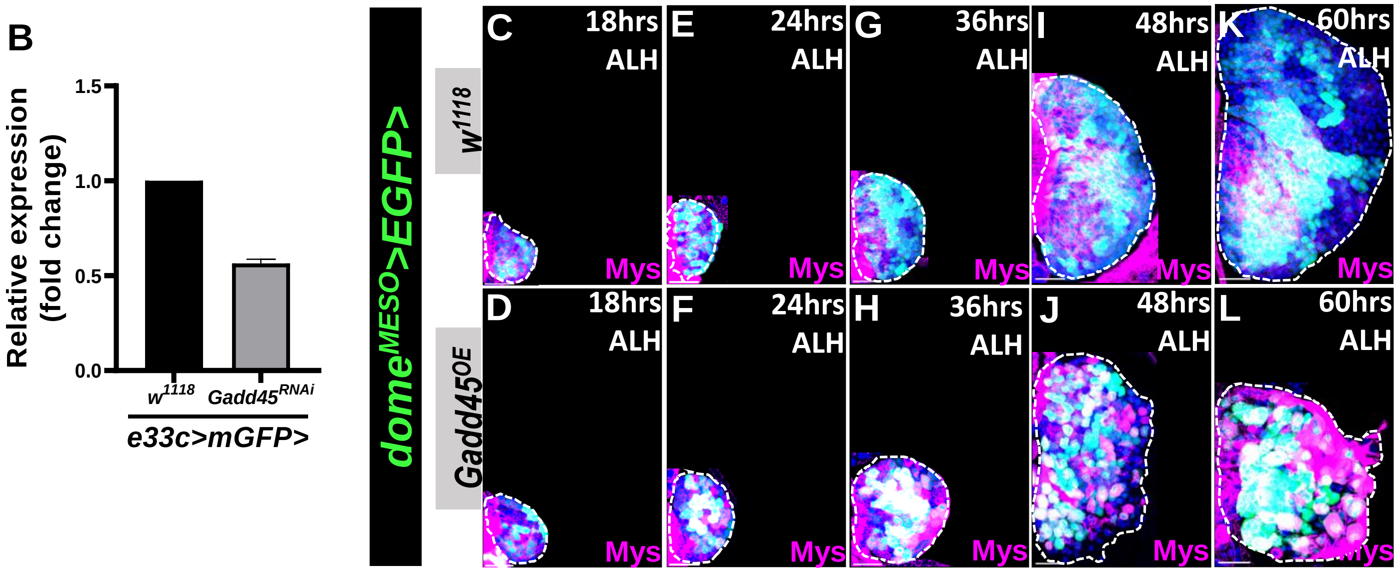


M

| Study | Cell populations/region | Key finding on Gadd45 expression |
|---|---|---|
| Cattenoz *et al.* (2020) | Embryonic and larval hemocytes | Gadd45 expression was higher in wandering L3 than in embryonic hemocytes. |
| Cho *et al.* (2020) | Lymph gland plasmatocytes, crystal cells and lamellocytes | Gadd45 transcripts were detected across multiple hemocyte populations, with variation among clusters. |
| Tattikota *et al.* (2020) | Larval circulating plasmatocyte and crystal-cell subclusters | Gadd45 expression varied among hemocyte subclusters. |
| Girard *et al.* (2021) | Lymph gland: intermediate-zone and plasmatocyte-associated populations | Gadd45 transcripts were enriched in intermediate-zone, proplasmatocyte-associated and plasmatocyte-associated populations. |

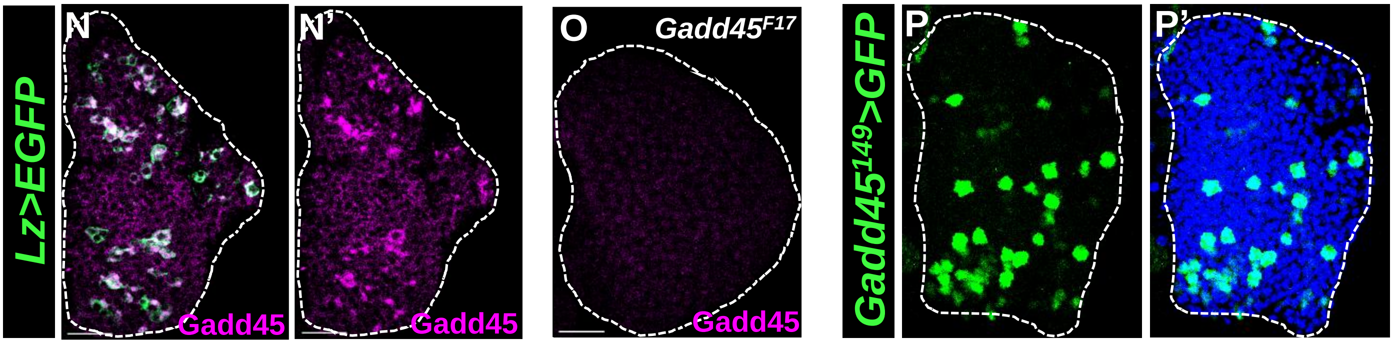

#### Figure S1: Gadd45 in third-instar lymph gland

**(A)** A 100 bp DNA ladder is used as a reference DNA marker. The lanes containing positive control DNA from *w*$^{1118}$ exhibited a single, distinct band with Rp49, F1/R1, and F1/R3 primers, confirming that the primers and reaction conditions were functional. No band appears in the *Gadd45*$^{F17}$ lane with primer pair F1/R1, while a smaller band is visible in the *Gadd45*$^{F17}$ lane with primer pair F1/R3, confirming the deletion mutation. NTC (No Template Control) lanes are included for each primer set.

**(B):** Quantitative RT-PCR analysis showing relative expression of *Gadd45* transcripts in lymph glands expressing *Gadd45*$^{RNAi}$ driven by *e33c>mGFP* compared to control *e33c>mGFP/+*. *Gadd45*$^{RNAi}$ results in significant reduction in *Gadd45* transcript levels.

**(C-L)** Merged images showing *dome*$^{MESO}$*>EGFP* (green) marking progenitors, Mys (magenta) marking lamellocytes, and DAPI (blue) marking nuclei to visualize lymph gland size. Gadd45 overexpression does not alter Mys expression at 18 h ALH (D, n = 4), comparable to control lymph glands *dome*$^{MESO}$*-Gal4, UAS-EGFP/+* (C, n = 4). At 24 h ALH, Gadd45 overexpression initiates lamellocyte differentiation (F, n = 14), whereas control retains Mys expression primarily in progenitor cells (E, n = 12). This progressively increases at 36 h ALH (H, n = 12) compared to control (G, n = 10) and results in extensive lamellocyte production by 48 h ALH (J, n = 12) compared to control (I, n = 10). By 60 h ALH, control lymph glands continue to exhibit progenitor-level Mys expression (K, n = 6), whereas Gadd45-overexpressing lymph glands show strong Mys upregulation, consistent with extensive lamellocyte differentiation and cortical zone disintegration (L, n = 6).

**(M)** Summary of published single-cell transcriptomic studies reporting Gadd45 expression across *Drosophila* hemocyte populations. The table summarizes findings from Cattenoz et al. (2020), Cho et al. (2020), Tattikota et al. (2020), and Girard et al. (2021), highlighting the developmental stages, hemocyte populations, and lymph-gland regions in which Gadd45 transcripts were detected or showed expression variation. Collectively, these studies indicate that Gadd45 expression varies across hemocyte populations and developmental stages, providing a transcriptomic basis for investigating its potential role in hematopoietic development and lineage regulation.

**(N-O)** Third instar lymph glands expressing *Lz>EGFP* (green, crystal cells) show Gadd45 transcripts (magenta) by RNA FISH. Gadd45 mRNA is detected in Lz+ crystal cells (N-N', n=20), whereas loss of Gadd45 FISH signal in the *Gadd45*$^{F17}$ null mutant lymph gland (O, n=16) confirms probe specificity.

**(P-P')** In the third instar, in the *Gadd45*$^{149}$*-Gal4 UAS-EGFP* background, a subset of cells, designated Gadd45>GFP+, is distributed throughout the lymph gland.

All confocal images are maximum-intensity projections of the middle third of the optical sections of the wandering third instar lymph gland lobe from at least three independent biological experiments. White dashed lines outline a lymph gland lobe. Scale bar represents 25 µm. DAPI (blue)-stained nuclei.

Figure S2

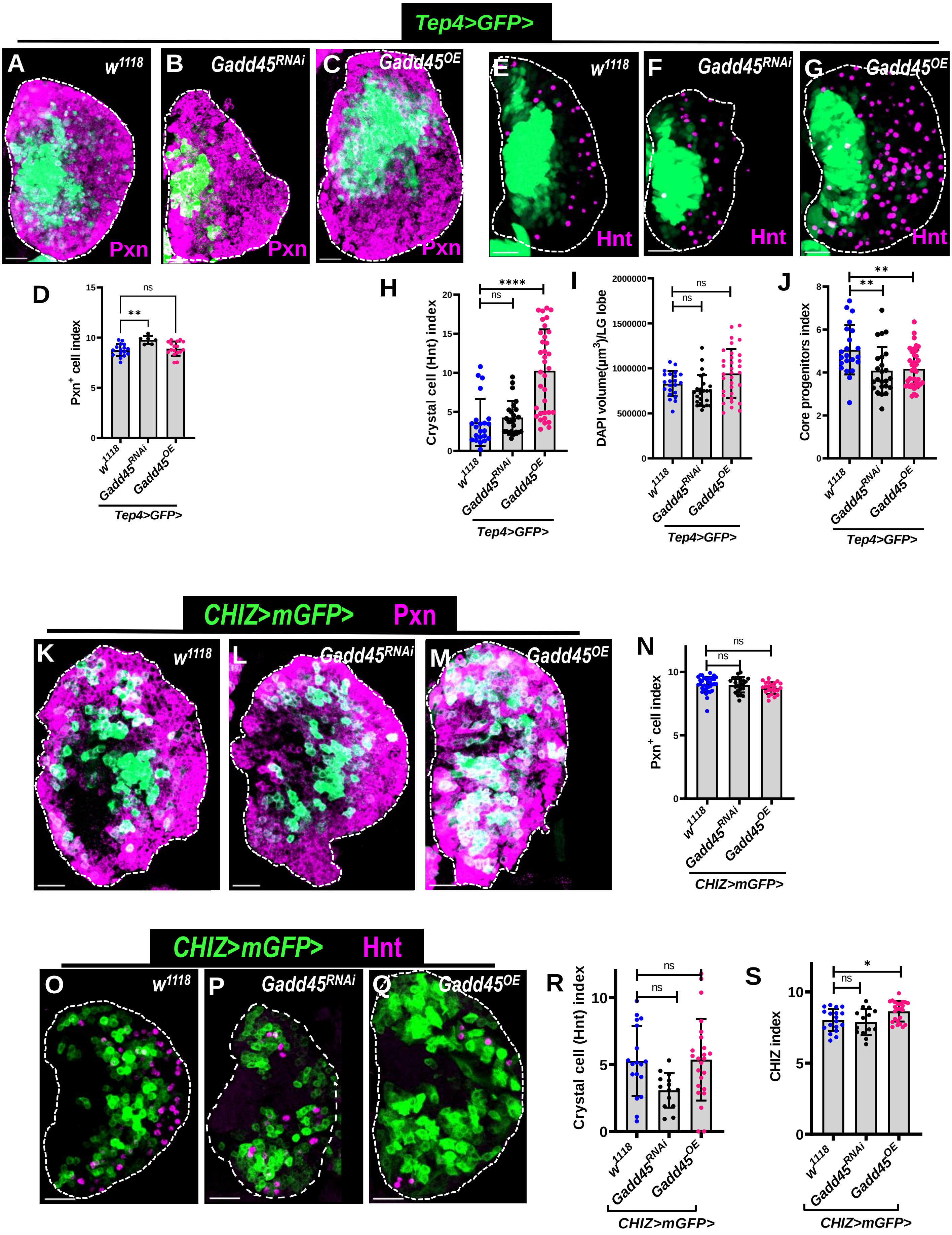

Tep4>GFP>
A w1118 Pxn
B Gadd45RNAi Pxn
C Gadd45OE Pxn
E w1118 Hnt
F Gadd45RNAi Hnt
G Gadd45OE Hnt
D
Pxn+ cell index
15
10
5
0
**
ns
w1118
Gadd45RNAi
Gadd45OE
Tep4>GFP>
H
Crystal cell (Hnt) index
20
15
10
5
0
****
ns
w1118
Gadd45RNAi
Gadd45OE
Tep4>GFP>
I
DAPI volume(μm3)/LG lobe
2000000
1500000
1000000
500000
0
ns
ns
w1118
Gadd45RNAi
Gadd45OE
Tep4>GFP>
J
Core progenitors index
8
6
4
2
0
**
**
w1118
Gadd45RNAi
Gadd45OE
Tep4>GFP>
CHIZ>mGFP> Pxn
K w1118
L Gadd45RNAi
M Gadd45OE
N
Pxn+ cell index
10
5
0
ns
ns
w1118
Gadd45RNAi
Gadd45OE
CHIZ>mGFP>
CHIZ>mGFP> Hnt
O w1118
P Gadd45RNAi
Q Gadd45OE
R
Crystal cell (Hnt) index
10
5
0
ns
ns
w1118
Gadd45RNAi
Gadd45OE
CHIZ>mGFP>
S
CHIZ index
10
5
0
*
ns
w1118
Gadd45RNAi
Gadd45OE
CHIZ>mGFP>

**Figure S2. Dysregulation of GADD45 in lymph gland intermediate progenitors causes lamellocytes differentiation.**

**(A-C)** Differential Gadd45 expression using *Tep4-Gal4, UAS-2XEGFP* (green) by expressing *Gadd45*$^{RNAi}$ (B, n=6) and *Gadd45*$^{OE}$ (C, n=17) does not show any visible alteration in the volume of Pxn+ cells (Pxn: peroxidasin, an early plasmatocyte maker) compared to control (A, n=16).

**(D)** Quantification of Pxn+ cell index in (A-C).

**(E-G)** Core progenitors specific *Tep4-Gal4, UAS-2XEGFP* (green) mediated *Gadd45*$^{RNAi}$ (F, n=24) expressing lymph glands do not show any change in crystal cells number, whereas *Gadd45*$^{OE}$ (G, n=35) expressing lymph glands have significantly increased number of crystal cells when compared with the control (*Tep4-Gal4, UAS-2XEGFP/+*) (E, n=22).

**(H)** Quantification of crystal cell index in (E-G).

**(I)** Quantification of lymph gland size using DAPI volume (E-G).

**(J)** Quantification of core progenitor index in (E-G).

**(K-M)** Expression of *UAS-Gadd45*$^{RNAi}$ (L, n=24) and *UAS-Gadd45*$^{OE}$ (M, n=24) in the Intermediate progenitors (green) using *CHIZ-Gal4, UAS-mGFP* driver, left the Pxn positive cell number (magenta) unchanged compared to the control (K, n=36)

**(N)** Quantification of Pxn+ cell index in (K-M).

**(O-Q)** Knockdown and expression of Gadd45 in intermediate progenitors (green) using *CHIZ-Gal4, UAS-mGFP* by expressing *UAS-Gadd45*$^{RNAi}$ (P, n=14) and *UAS-Gadd45*$^{OE}$ (Q, n=24) did not show any significant difference in crystal cell numbers marked by Hnt (magenta) compared to control (O, n=18) lymph glands.

**(R)** Quantification of crystal cells index in (O-Q).

**(S)** Quantification of CHIZ (intermediate progenitor) index in (O-Q).

All confocal images are maximum-intensity projections of the middle third of the optical sections of the wandering third instar lymph gland lobe from at least three independent biological experiments. White dashed lines outline a lymph gland lobe. Scale bar represents 25 µm. Error bars in graphs: mean ± S.D. ns.: not significant (p>0.01); *p<0.01; **p<0.001; ****p<0.00001. n represents lymph gland lobes.

Figure S3

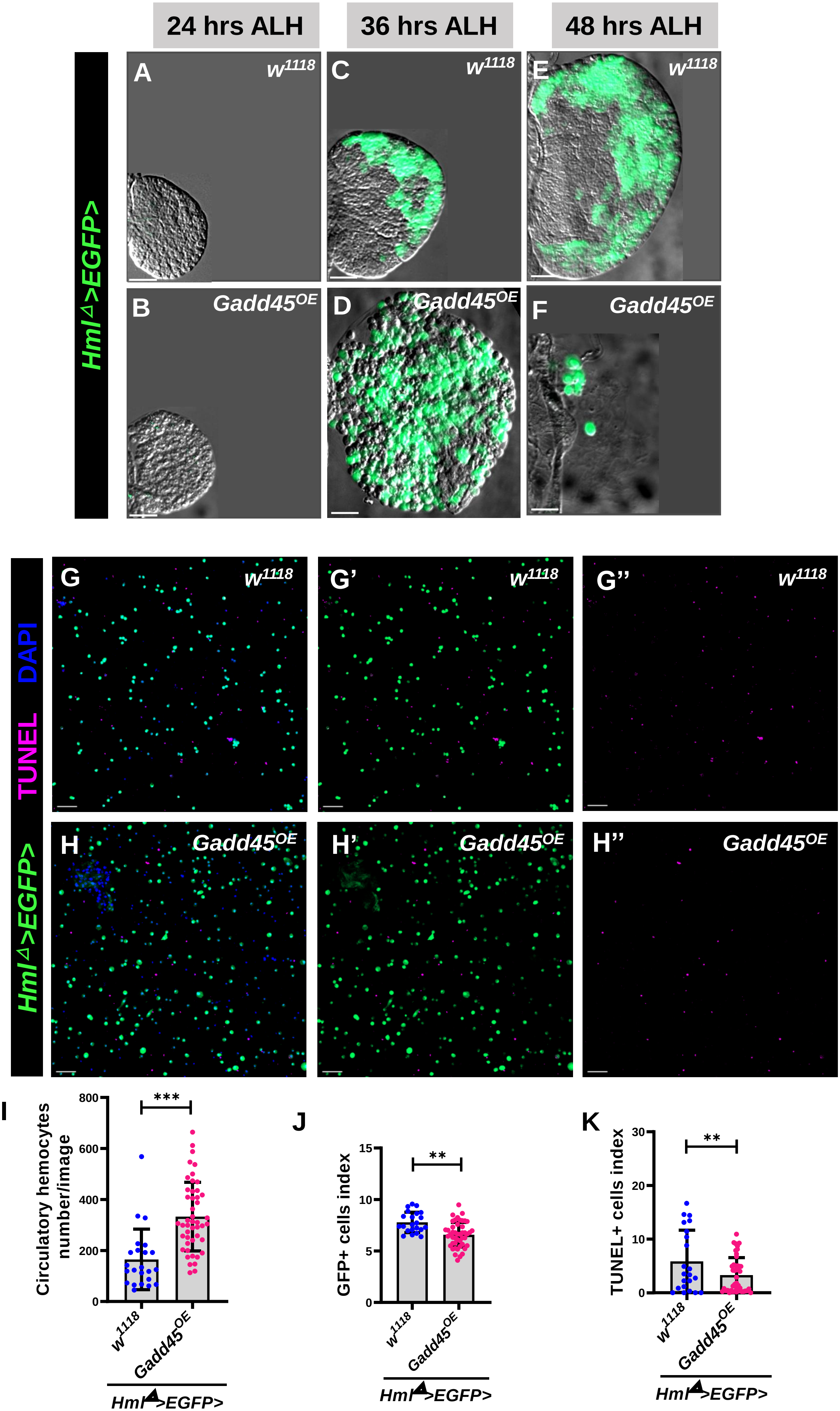

24 hrs ALH
36 hrs ALH
48 hrs ALH
HmlΔ>EGFP>
A
w1118
B
Gadd45OE
C
w1118
D
Gadd45OE
E
w1118
F
Gadd45OE
HmlΔ>EGFP> TUNEL DAPI
G
w1118
G'
w1118
G''
w1118
H
Gadd45OE
H'
Gadd45OE
H''
Gadd45OE
I
Circulatory hemocytes number/image
800
600
400
200
0
***
w1118
Gadd45OE
HmlΔ>EGFP>
J
GFP+ cells index
15
10
5
0
**
w1118
Gadd45OE
HmlΔ>EGFP>
K
TUNEL+ cells index
30
20
10
0
**
w1118
Gadd45OE
HmlΔ>EGFP>

**Figure S3. Gadd45 overexpression in hemocytes causes premature lymph gland disintegration without increasing cell death**

**(A-F)** An early differentiation zone marker, *Hml*$^{\Delta}$*-Gal4*-driven *Gadd45* overexpression at 24 h ALH; *Gadd45*-overexpressing lymph gland primary lobes (B, n=18) were comparable in overall appearance to controls (A, n=34). At 36 h ALH, the primary lobes of *Gadd45*-overexpressing larvae (D, n=20) showed a pronounced increase in size compared with controls (C, n=30). By 48 h ALH, the lymph gland lobes in *Gadd45*-overexpressing larvae were largely disintegrated (F, n=24), whereas the control lymph glands remained intact (E, n=40).

**(G-H")** Circulatory hemocytes from control *Hml*$^{\Delta}$*>EGFP/+* (G-G") (n=24) and *Hml*$^{\Delta}$*>EGFP>Gadd45*$^{OE}$ (H-H") (n=44) third-instar larvae stained for apoptosis. The GFP+ and TUNEL+ cell indices are reduced upon Gadd45 overexpression, indicating no induction of apoptosis, whereas the total number of circulating hemocytes is significantly increased in *Gadd45*$^{OE}$ compared to control.

**(I)** Quantification of circulating hemocytes per larvae in (G-H'').

**(J)** Quantification of circulating GFP positive cell index in (G-H'').

**(K)** Quantification of circulating TUNEL positive cell index in (G-H'').

All confocal images represent maximum-intensity projections of wandering third-instar circulating hemocytes from at least three independent biological experiments. Scale bar represents 50 µm. DAPI (blue)-stained nuclei. Error bars in graphs: mean ± S.D. ns.: not significant (p>0.01); **p<0.001; ****p<0.00001. n represents the number of larvae or lymph gland lobes.

**Figure S4**

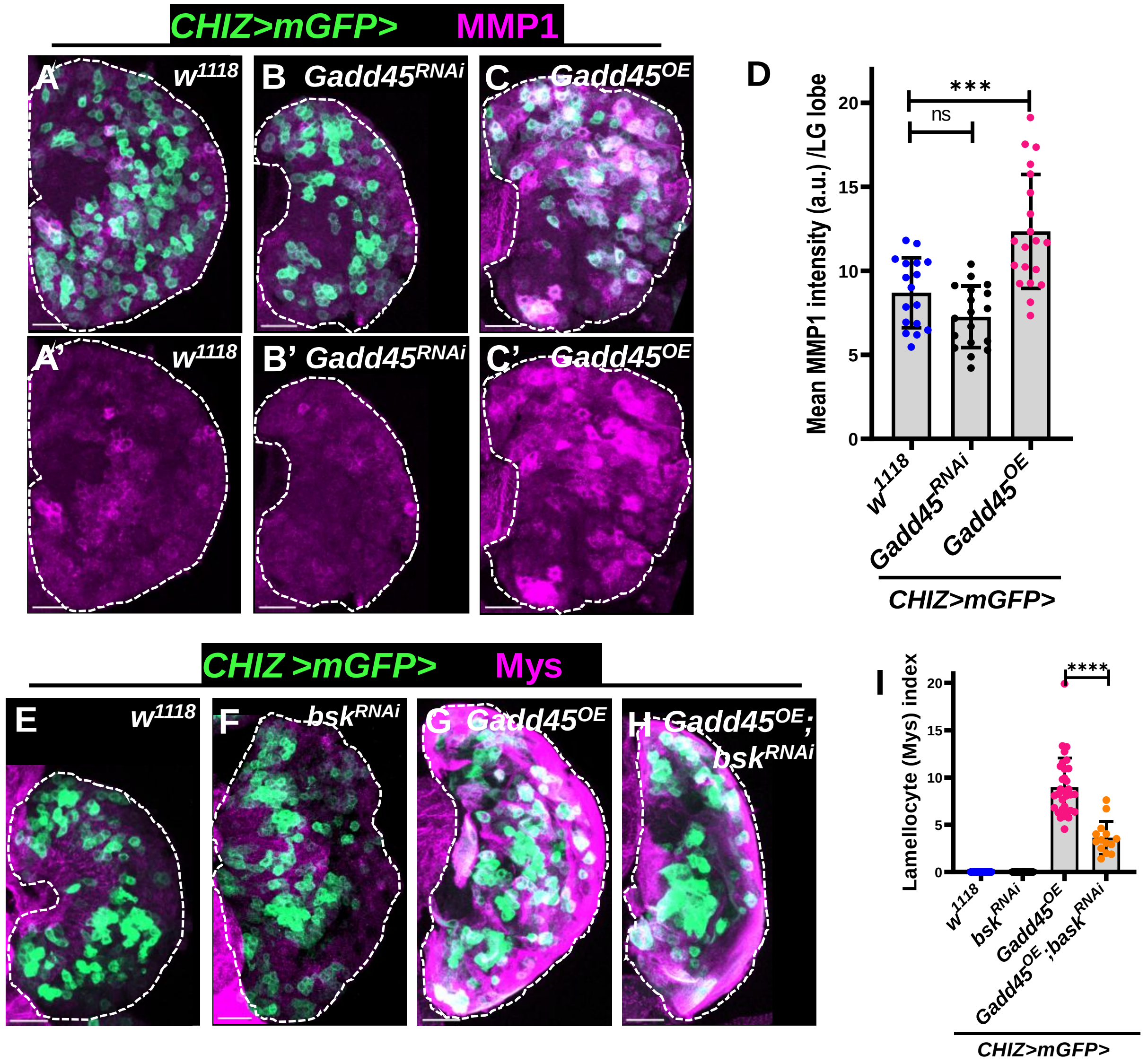


**Figure S4. Gadd45 overexpression activates JNK signaling, crucial for lamellocyte differentiation.**

**(A-C')** Matrix metalloprotease 1 **(**MMP1) immunostaining (magenta) shows JNK signaling activity, in *CHIZ>mGFP* (green)-driven *Gadd45*$^{RNAi}$ (B-B', n=20) resulted in the same MMP1 expression (magenta) as the control (A-A', n=20), while in *Gadd45*$^{OE}$ (C-C', n=20), it increased significantly.

**(D)** Quantification of mean fluorescent intensity of MMP1 per lymph gland lobe in (A-C').

**(E-H)** Intermediate progenitor-specific knockdown of *bsk*$^{RNAi}$ (F, n=13) does not induce lamellocyte differentiation, resembling the control (*CHIZ-Gal4, UAS-mCD8::GFP/+*) (E, n=10) lymph glands. Expression of *Gadd45*$^{OE}$ in the background of *bsk*$^{RNAi}$ within the intermediate progenitor (green) significantly decreases the *Gadd45*$^{OE}$-mediated lamellocytes number (J, n=14) compared to the lymph glands expressing *Gadd45*$^{OE}$ (H, n=12).

**(I)** Quantification of lamellocytes index in (E-H).

All confocal images are maximum-intensity projections of the middle third of optical sections of the wandering third-instar lymph gland lobe from at least three independent biological experiments. White dashed lines outline a lymph gland lobe. Scale bar represents 25 µm. Error bars in graphs: mean ± S.D. ns.: not significant ($p>0.01$); ***$p<0.0001$; ****$p<0.00001$. n represents lymph gland lobes.

### Figure S5

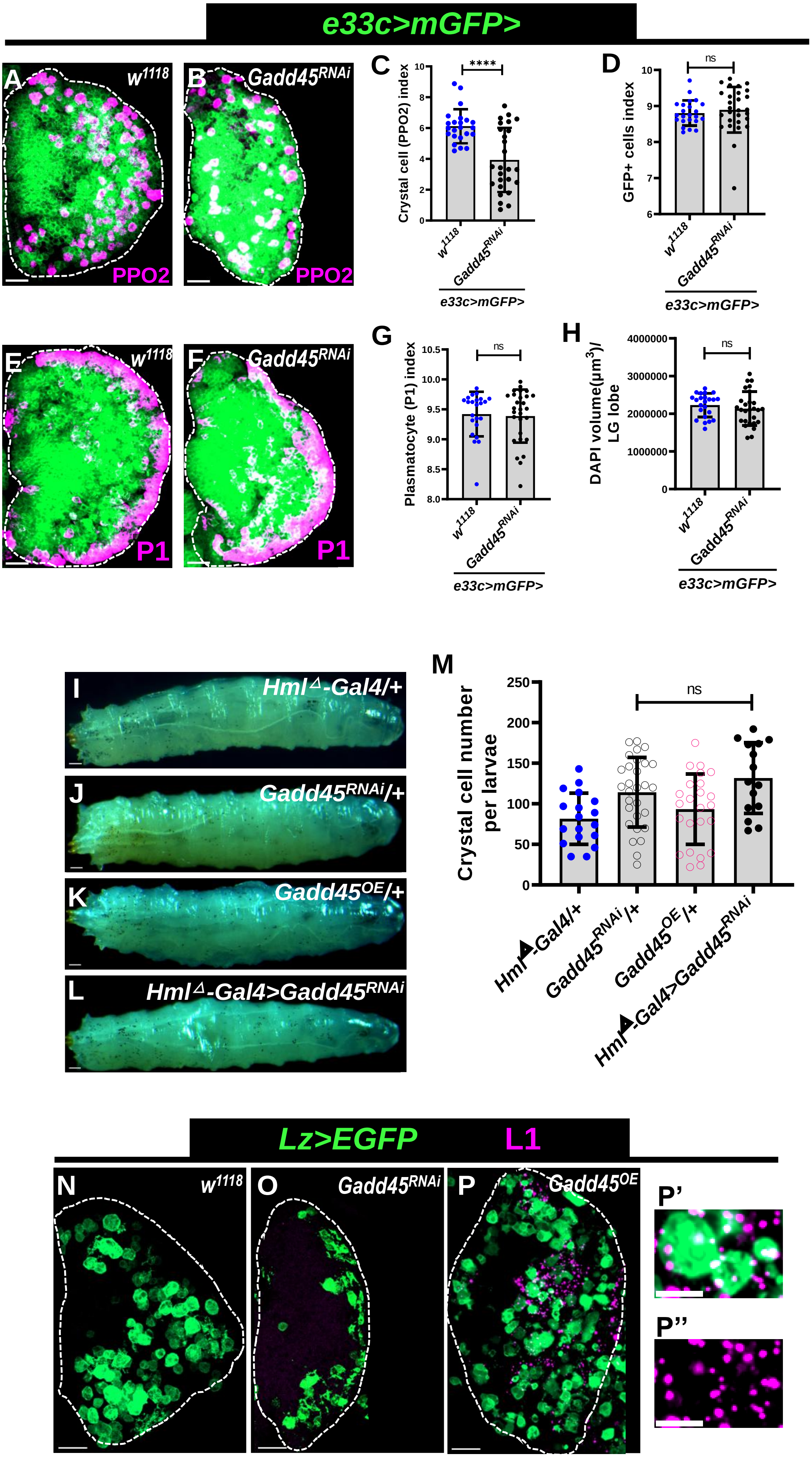

e33c>mGFP>
A
w1118
PPO2
B
Gadd45RNAi
PPO2
C
Crystal cell (PPO2) index
****
w1118
Gadd45RNAi
e33c>mGFP>
D
GFP+ cells index
ns
w1118
Gadd45RNAi
e33c>mGFP>
E
w1118
P1
F
Gadd45RNAi
P1
G
Plasmatocyte (P1) index
ns
w1118
Gadd45RNAi
e33c>mGFP>
H
DAPI volume(µm3)/ LG lobe
ns
w1118
Gadd45RNAi
e33c>mGFP>
I
HmlΔ-Gal4/+
J
Gadd45RNAi/+
K
Gadd45OE/+
L
HmlΔ-Gal4>Gadd45RNAi
M
Crystal cell number per larvae
ns
HmlΔ-Gal4/+
Gadd45RNAi/+
Gadd45OE/+
HmlΔ-Gal4>Gadd45RNAi
Lz>EGFP
L1
N
w1118
O
Gadd45RNAi
P
Gadd45OE
P'
P''

#### Figure S5. Loss of Gadd45 impairs crystal cell formation but not plasmatocyte differentiation

**(A-B):** Gadd45 was knocked down using a pan-lymph gland driver *e33c-Gal4, UAS-mCD8-GFP* and immunostained for crystal cell marker PPO2 (magenta). *e33c>mGFP>Gadd45*$^{RNAi}$ (B, n=30) lymph glands show a drastic decrease in PPO2+ crystal cells compared to control *e33c>mGFP/+* (A, n=23).
**(C):** Quantification of crystal cell (PPO2) index in (A-B).
**(D):** Quantification of GFP+ cell index (A-B).
**(E-F):** Gadd45 depletion using a pan-lymph gland driver *e33c>mGFP>Gadd45*$^{RNAi}$ (F, n=30) does not alter the number of mature plasmatocytes marked by P1 (magenta) in comparison to control lymph glands *e33c>mGFP/+* (E, n=23).
**(G):** Quantification of plasmatocyte (P1) index in (E-F).
**(H):** Quantification of lymph gland size using DAPI volume in (E-F).
**(I-L)**: Wandering third-instar larvae of the indicated genotypes were heat-shocked (60°C, 10 min) to induce crystal cell melanization, visible as black body-wall spots. Spot numbers were comparable in controls *HmlΔ-Gal4/+* (I, n=18), *Gadd45*$^{RNAi}$*/+* (J, n=28), *Gadd45*$^{OE}$*/+* (K, n=23), and larvae with differentiated hemocyte-specific Gadd45 knockdown, *HmlΔ-Gal4>Gadd45*$^{RNAi}$ (L, n=14).
**(M)** Quantification of spots per larva in (I-L).
**(N-P”)** Crystal cells are marked by *Lz-Gal4, UAS-EGFP* (green) and lamellocytes by L1 (magenta). Control *Lz>EGFP/+* (N, n=10) and *Lz-Gal4, UAS-EGFP*-driven *Gadd45*$^{RNAi}$ (O, n=14) lymph glands show *EGFP+* crystal cells negative for L1 whereas *Lz-Gal4, UAS-EGFP*-driven *Gadd45*$^{OE}$ (P, n=14) shows ectopic induction of L1 within *Lz+* crystal cells. The boxed area in (P) is magnified in (P', P''), showing colocalization of L1 (magenta puncta) inside *EGFP*-positive crystal cells.

All confocal images represent maximum-intensity projections of the middle-third optical sections, except (K', K''), which is a single optical section of the wandering third-instar lymph gland lobe. White dashed lines outline a lymph gland lobe. Scale bar represents 25 µm. Error bars in graphs: mean ± S.D. ns.: not significant (p>0.01); ***p<0.0001. n represents lymph gland lobes.

#### Figure S6

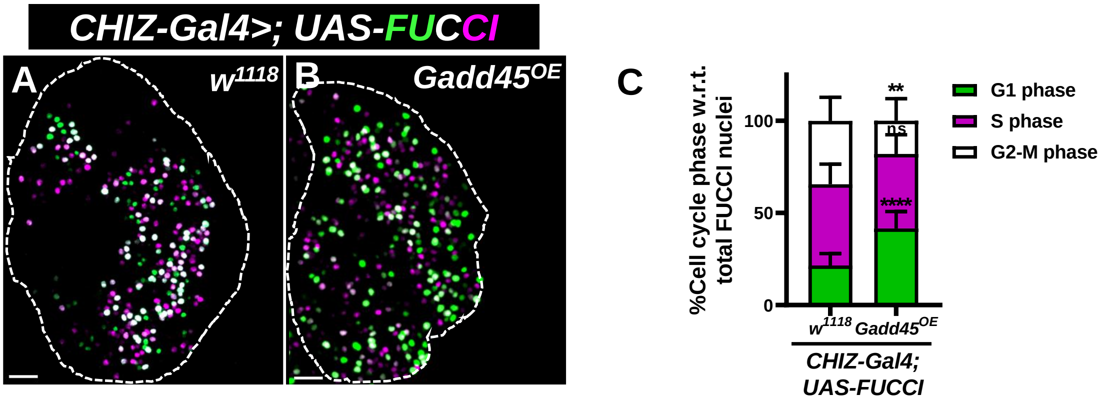


**Figure S6. Gadd45 overexpression causes G1 arrest in lymph gland progenitors**

**(A-B)** Intermediate progenitor-specific expression of *Gadd45$^{OE}$* (*CHIZ-Gal4, UAS-FUCCI*-driven *UAS-Gadd45$^{OE}$)* (B, n=19) induces G1-phase (green only) arrest compared to control (A, n=11).

**(C)** Quantification of %G1, %S, and %G2/M cells in (A-B).

All confocal images represent maximum-intensity projections of the middle third optical sections of the wandering third-instar lymph gland lobe. White dashed lines outline a lymph gland lobe. Scale bar represents 25 µm. Error bars in graphs: mean ± S.D. ns.: not significant (p>0.01); ***p<0.0001. n represents lymph gland lobe numbers.

Figure S7

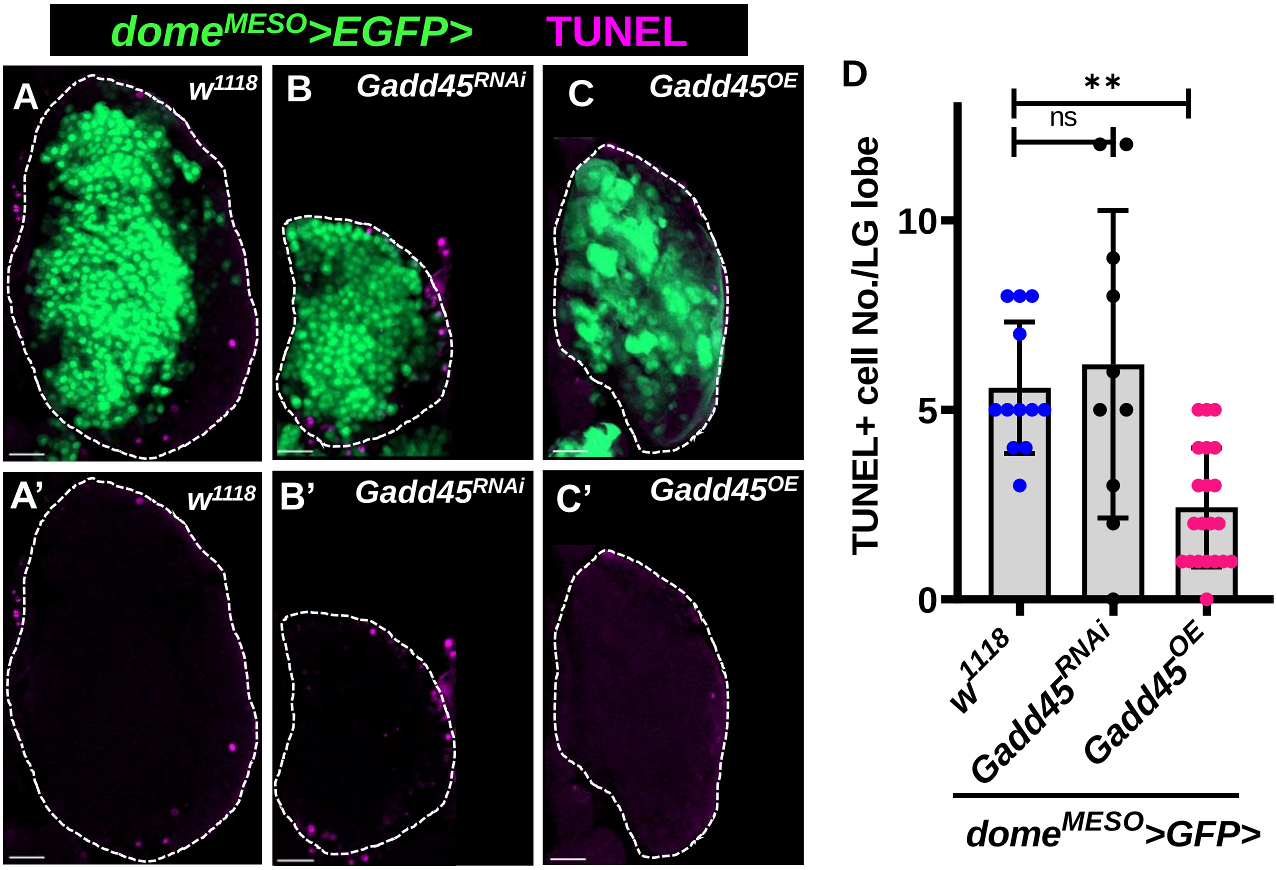


#### Figure S7. No impact on apoptosis upon Gadd45 dysregulation in lymph gland progenitors

**(A-C')** Lymph gland progenitors specific *dome*$^{MESO}$*-Gal4, UAS-2xEGFP* driven *UAS-Gadd4*$^{RNAi}$ (B-B', n=10) show a similar number of TUNEL-positive (magenta) cells. However, *UAS-Gadd45*$^{OE}$ (C-C', n=21) shows fewer TUNEL-positive (magenta) cells than the control (*dome*$^{MESO}$*-Gal4, UAS-2xEGFP*/+) lymph glands (A-A', n=12).

**(D)** Quantification of TUNEL-positive (apoptotic marker) cells per anterior lymph gland lobe in (A-C').

All confocal images represent maximum-intensity projections of the middle-third optical sections of the wandering third-instar lymph gland lobe. White dashed lines outline a lymph gland lobe. Scale bar represents 25 μm. Error bars in graphs: mean ± S.D. ns.: not significant (p>0.01); ***p<0.0001. n represents lymph gland lobes.
